# HyRes: Accurate Physics-Based Simulation of Dynamic Protein Structures and Interactions in Complex Environments at Scale

**DOI:** 10.64898/2026.06.23.734133

**Authors:** Shanlong Li, Shrishti Barethiya, Jianhan Chen

## Abstract

Intrinsically disordered proteins and regions (IDPs) are ubiquitous cellular regulators. Uncovering how their transient, multivalent interactions organize and fine-tune cellular processes requires transferable methods capable of deriving dynamic conformational ensembles across diverse environments at scale. Here, we present HyRes, a physics-based, hybrid-resolution protein model with atomistic backbones and intermediate-resolution sidechains that bridges the gap between atomistic accuracy and computational efficiency. Evaluated across ∼100 IDPs, HyRes generates atomistic ensembles that match or outperform state-of-the-art all-atom force fields in reproducing experimental chain dimensions, transient tertiary contacts, and local secondary structures. Demonstrating exceptional transferability, HyRes accurately captures dynamic IDP interactions in dilute phases, condensed phases, and amyloid fibril fuzzy coats. Finally, we leverage HyRes’ scalability to generate disordered ensembles for ∼30,000 IDPs from the human proteome and DisProt, revealing strong correlation between residual structures and cellular function and localization. HyRes and this open-access database provide unprecedented resources for IDP biology and deep learning.

## Introduction

Cellular function relies on remarkable network plasticity and dynamic adaptability to respond to signals, stresses, and environmental cues^1,2^. This versatility is largely driven by highly flexible molecules, particularly intrinsically disordered proteins and proteins with long intrinsically disordered regions (IDPs)^3–5^. IDPs account for >30% of eukaryotic proteins and are central components of the regulatory networks that dictate virtually all aspects of cellular decision-making^6–8^. IDPs often remain unstructured even in specific complexes and functional assemblies^9–11^, highlighting a dynamic mode of specific protein interaction that is widespread but understudied^12^. Furthermore, these multivalent biomolecules are the major drivers of phase separation, the defining mechanism for the formation of membraneless organelles and an area of intense current interests^13–16^. Critically, dysregulation of IDPs contributes to many human diseases, including cancers, diabetes, neurodegenerative diseases, and heart diseases^17–19^.

Resolving the dynamic protein conformational ensembles in diverse environments is critical towards establishing the molecular mechanisms of IDP function and regulation^20^. How the disordered ensemble responds to various stimuli encodes function, whether it is cooperative folding upon binding or conformational redistribution in dynamic interactions^21^. Advances in nuclear magnetic resonance (NMR) spectroscopy^22^, small-angle scattering (SAS)^23^, and single-molecule Forster resonance energy transfer (smFRET)^24^, as well as other complementary techniques^25,26^, have significantly improved the ability to characterize disordered conformational ensembles at up to residue resolution. However, these measurements almost exclusively report on ensemble-averaged properties; it is a fundamentally underdetermined problem to resolve disordered ensembles using these experimental data alone^27,28^. Computational methods are thus required for generating full conformational ensembles of IDPs, particularly physics-based molecular simulation approaches^29–32^. The need for reliable simulation of disordered protein ensembles has actually been a major driver behind significant improvements in the state-of-the-art all-atom protein force fields in recent years, notably AMBER03ws (a03ws)^33^, CHARMM36m (C36m)^34^, AMBER99SB-disp (a99SB-disp)^35^, and others. Note that an important advantage of physics-based simulations is the versatility and transferability. In principle, they can be applied to probe the effects of almost any physical forces, chemical modifications, molecular interactions, and/or heterogeneous environments on IDP structure, dynamics, and function.

Atomistic simulations of IDPs in explicit solvent are computationally demanding due to the need to sample the broad manifold of relevant conformational space. Standard molecular dynamics (MD) is rarely adequate to generate converged IDP ensembles, even with tens of microseconds of sampling on ANTON supercomputers^35^; enhanced sampling such as replica exchange is generally required for achieving sufficient convergence to resolve the effects of mutation or binding^36,37^. The prohibitive cost of atomistic simulations has motivated extensive efforts to develop more efficient physics-based protein models with reduced resolutions, including implicit solvent^38–40^ and coarse-grained (CG) models^31,41–45^. At present, only one-bead-per-residue (*i.e.*, Cα-only) models have demonstrated successes in proteome-scale investigations of IDP sequence-ensemble–function relationships^30^. A critical challenge of these CG models is to capture sequence-dependent residual local and long-range structures that can play central roles in IDPs’ ability to support specific functions in cellular signaling and regulation^3,12,46^. Generative machine-learning (ML) models have also been trained using outputs from physics-based simulations to directly generate disordered ensembles from sequences with various levels of success^47–50^. These ML models offer efficient surrogates for the original simulation approaches, but they are strongly constrained by the training conditions and generally can’t probe how the disordered ensemble may respond to binding, chemical modifications, and other cellular signals for studying IDP function and regulation.

In this work, we address the critical limitations in computing dynamic protein conformational ensembles by developing a well-balanced, physics-based hybrid-resolution (HyRes) protein force field that can accurately simulate IDP structures and interactions across diverse environments at scale. HyRes combines atomistic backbones with intermediate-resolution side chains^51,52^ (**Fig. 1a**) to accurately describe backbone-mediated interactions and transient local and global structures. Early versions of HyRes have demonstrated a strong potential for studying IDP structure and interaction in solution, in complexes, and in condensates^53–56^. The HyRes force field is refined using radius of gyration (*R*_g_) measurements on 20 variants of the human hnRNPA1 low-complexity domain (A1-LCD) and NMR-derived residue helicity of 15 IDPs. Cross-validated using approximately 100 IDPs with various experimental data on both local and global structural properties, HyRes generates atomistic ensembles that are comparable, and often superior, to those derived from the state-of-the-art all-atom explicit solvent protein force fields, but with several orders of magnitude lower computational costs. We further demonstrate that HyRes can accurately describe dynamic IDP interactions in dilute and condensed phases, as well as in the fuzzy coats of amyloid fibrils, highlighting its outstanding transferability. Leveraging HyRes’ accuracy and efficiency, we further generate detailed conformational ensembles for over 30,000 IDPs in the human proteome^30^ and the DisProt database^4^. Together, HyRes and the proteome-scale open-access database of disordered conformational ensembles provide highly valuable resources for studying IDP structure and function and for enabling deep learning in IDP biology.

**Fig. 1.**
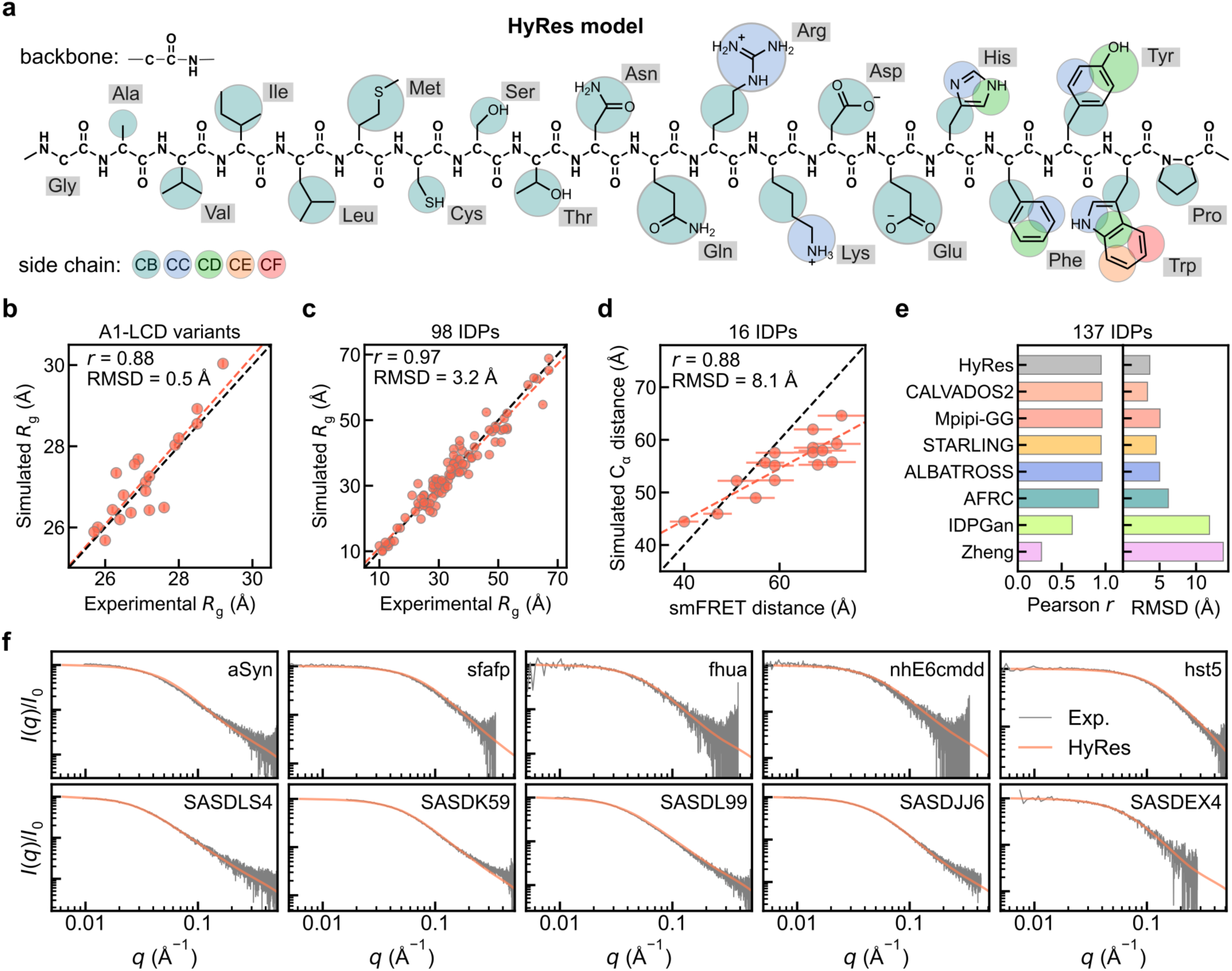
HyRes representation and prediction of IDP dimension. **a,** Schematic of residue representation in HyRes, featuring an atomistic backbone, -C-(CO)-(NH)-, and coarse-grained side chains (color-coded CB-CF beads). **b, c,** Simulated and experimental *R*g values for 20 A1-LCD variants (training) (**b**) and 98 additional IDPs with high-quality experimental data available (test) (**c**) (also see **Supplementary Tables S1 and S3**). **d,** Average end-to-end Cα distances of HyRes ensembles in comparison to experimental dye-pair distances derived from smFRET for 16 IDPs^49^ (also see **Supplementary Table S4**). In **b-d**, error bars plot standard deviations from block averaging analysis. **e,** Performance of HyRes relative to several state-of-the-art ML and CG models in predicting *R*g for 137 IDPs, as reported in Novak et al., Nature (2026)^49^. **f,** Representative SAXS scattering profiles calculated from HyRes ensembles in comparison with actual experimental measurements. Corresponding SASBDB entry IDs are noted.

## Results

### Rebalancing nonbonded interactions: accurate prediction of IDP chain dimension

To refine HyRes, electrostatic interactions were rebalanced through recalibration of the dielectric constant using the radii of gyration (*R*_g_) of glutamic acid-lysine (EK) peptides^57,58^, followed by refinement of backbone and side chain van der Waals (vdW) interactions against deca-arginine (R10)^59^ as well as 20 A1-LCD variants^60^ (see **Methods**, **Extended Data Fig. 1a-d,** and **Supplementary Tables S1-S2**). Note that the A1-LCD set contains perturbations to cation-π, π-π, hydrophobic, polar, and electrostatic contributions, thus providing a particularly effective basis for force field rebalancing^61^. As shown in **Fig. 1b**, the rebalanced parameter set achieves a small root-mean-square deviation (RMSD) of 0.5 Å for *R*_g_ of A1-LCD variants with a strong correlation of *r* = 0.88, which is very challenging because of the narrow range of ∼4 Å. Detailed examination of simulation-derived scattering profiles confirms that they closely match experimental SAXS measurements for all variants (**Extended Data Fig. 1f**), supporting that the conformational ensembles are accurately captured.

The refined model was critically evaluated using a large set of 98 IDPs spanning sequence lengths of 16 to 477 residues and *R*_g_ values of 10 to 70 Å (**Supplementary Table S3**). Across this set, the HyRes model achieves a high correlation of *r* = 0.97 with an RMSD of 3.2 Å (**Fig. 1c**). We further compare the average end-to-end distances (*R*_ee_) for 16 IDPs to smFRET-derived values^62^ (**Supplementary Table S4**), which achieves a correlation of *r* = 0.88 but with a higher RMSD of 8.1 Å (**Fig. 1d** and **Extended Data Fig. 1e**). The larger RMSD is expected because the dye molecules were not explicitly represented in HyRes simulations. We further confirmed that SAXS profiles derived from HyRes ensembles closely matched the experimental measurements for a curated set of 45 IDPs with available high-quality SAXS data^49^ (**Fig. 1f** and **Extended Data Fig. 2)**, indicating that HyRes captures both ensemble-averaged chain dimensions and underlying conformational distributions. Collectively, these results demonstrate that the refined HyRes model robustly reproduces various dimensional properties of IDPs with diverse sequences and levels of compactness. Importantly, HyRes’s ability to capture IDP chain dimensions is comparable to the best machine-learning predictors^49^ and data-driven CG models^61^ that have been explicitly trained on reproducing experimental *R*_g_ of larger numbers of IDPs (**Fig. 1e**).

### Ability of HyRes in capturing transient long-range interactions

IDPs frequently form transient long-range interactions that shape the conformational ensembles and modulate function^20^. To evaluate the ability of HyRes to capture such long-range structural properties, we collected paramagnetic relaxation enhancement (PRE) NMR data for a set of 9 IDPs with multiple spin-labeling sites to complement the above benchmarks on the global chain dimension (**Supplementary Table S5**). PRE is arguably the most powerful experimental approach for comprehensive analysis of transient long-range contacts within heterogeneous ensembles, requiring only a small number of strategic site-directed paramagnetic spin labeling constructs^64^. We used DEER-PREdict^65^ to calculate predicted PRE profiles from HyRes ensembles (see **Methods**). As summarized in **Fig. 2a** and **Extended Data Fig. 3**, HyRes successfully captures virtually all major features of the experimental PRE profiles for all 9 IDPs. For example, HyRes faithfully predicts the local compaction of protein osteopontin within residues 50-160, including pronounced contacts between residues 50-75 and 100-125 as revealed in the PRE profile with site 64 spin-labeled (**Fig. 2a**). Similar observations can be made for long-range contacts revealed in PRE profiles of protein RNAPd (**Extended Data Fig. 3d**). Across the set of 9 IDPs, HyRes reproduces experimental NMR PRE profiles with an average correlation of *r* ∼ 0.72 and average RMSD of 0.19 (**Fig. 2b** and **Supplementary Table S5**). Note that such a performance is similar to or better than that of ensembles generated from extensive simulations using several all-atom protein force fields including a99SB-disp, a03ws and C36m (**Fig. 2c** and **Supplementary Table S6**). The ability of HyRes to accurately capture these nontrivial long-range structural features of disordered ensembles is noteworthy, highlighting the strong balance that HyRes achieves across diverse physical interactions and sequences.

**Fig. 2.**
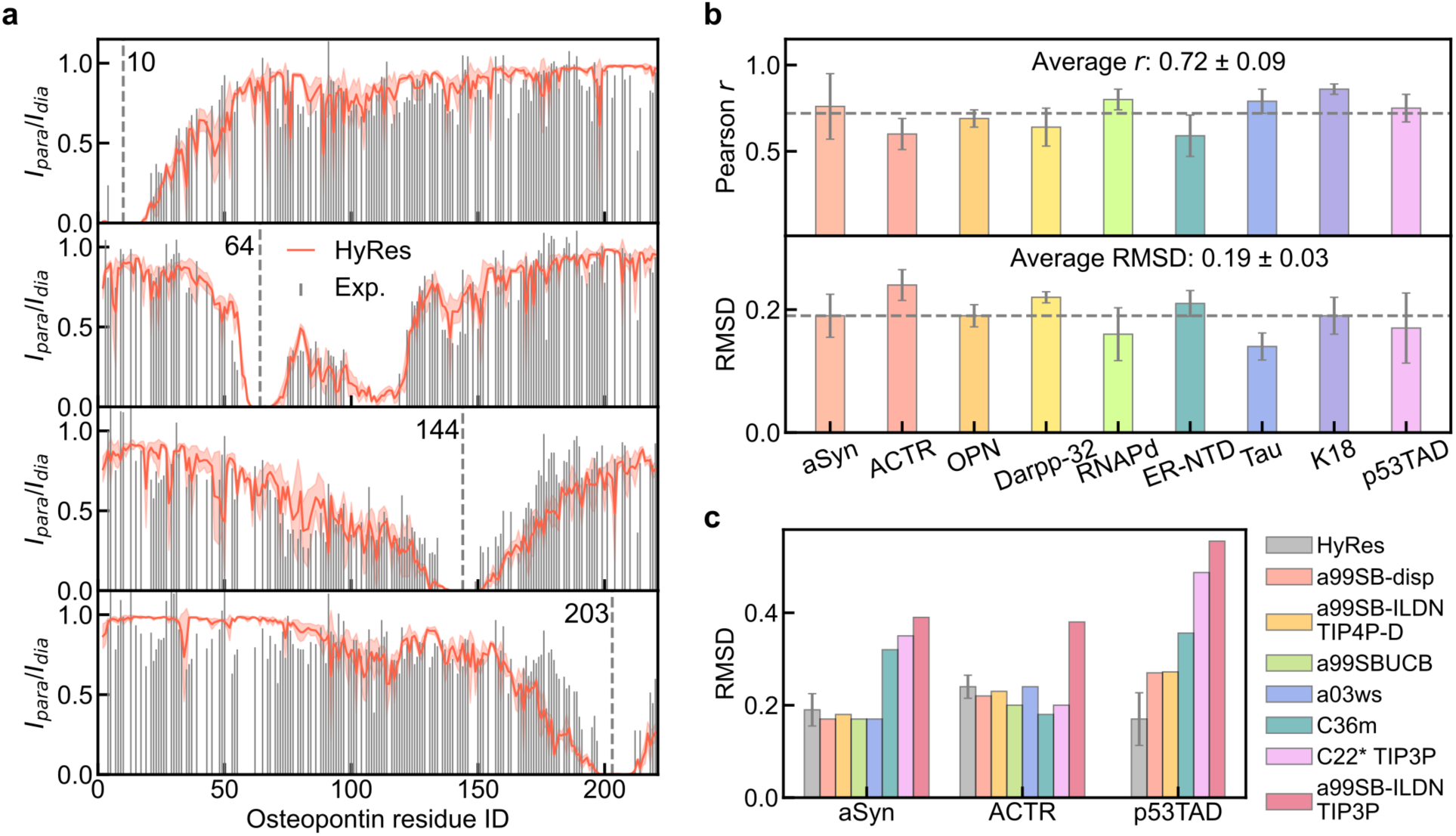
PRE benchmark of HyRes prediction of transient long-range contacts. **a,** PRE profiles derived from HyRes simulations (red lines) in comparison with experimental measurements (grey bars) for protein osteopontin. Spin-labeling sites are indicated by vertical dashed lines and residue numbers. Shaded pink regions represent standard deviations estimated from three independent HyRes simulations. **b,** Overall performance of HyRes in predicting PRE profiles across 9 IDPs. Error bars denote the standard deviations obtained from different spin-label positions. Grey dashed lines indicate average values. **c,** Performance of HyRes and representative all-atom protein force fields in predicting PRE profiles of aSyn^35^, ACTR^35^, and p53TAD^63^.

### Atomistic ensembles from HyRes: transient secondary structures of IDPs

Armed with an atomistic backbone, HyRes has a unique ability among CG models to capture transient secondary structures, which can be directly compared with residue-level propensities estimated from NMR secondary chemical shift analysis^66^. All-atom side chains can also be readily reconstructed with minimal computational cost, through backmapping or using ML tools such as cg2all^67^ with sub-Å accuracy. The resulting atomistic ensembles enable the estimation of NMR chemical shifts using SHIFTX2^68^ for direct comparison with experimental values. For this, we curated a set of 58 IDPs with available chemical shift assignments from the BMRB database^69^ (see **Methods**; **Supplementary Table S7**), with length ranging from 18 to 406 residues and average helicity ranging from approximately 0.0 to 0.3.

The backbone hydrogen bond strength was first calibrated using (AAQAA)_3_, followed by fine-tuning of helical propensity for each amino acid using a training set of 15 IDPs (**Fig. 3a**, **Extended Data Fig. 4a** and **Supplementary Table S8**). Residual helicities were calculated from HyRes trajectories using DSSP^70^ and compared with secondary structure propensity (SSP) scores^66^ derived from NMR chemical shifts or collected from published studies (**Fig. 3** and **Extended Data Fig. 4**). Across a large independent test set of 40 IDPs, the refined HyRes achieved a correlation of 0.83 and an average RMSD of 0.05 in mean helicity (**Fig. 3b** and **Supplementary Table S9**). There appears to be a tendency for HyRes to predict small and lightly populated helices absent from NMR secondary chemical analysis (e.g., **Fig. 3d** TC-1 N-terminal region), leading to slight bias towards overestimation for sequences with low (*e.g.*, <0.1) average helicity (**Fig. 3b**). We note that SSP scores can adopt both negative and positive values on a per-residue basis. Residues with negative SSP scores were set to zero when calculating experimental mean helicity, which may lead to underestimation of the experimentally inferred helicity. Importantly, HyRes successfully captured the effects of single mutations on average helicity of TDP-43 with a correlation of 0.9 (**Fig. 3c, Extended Data Fig. 6a** and **Supplementary Table S10**), reflecting a major strength of physics-based simulation rarely achievable with machine learning. Furthermore, HyRes correctly identifies most major helical segments (**Fig. 3d** and **Extended Data Fig. 4)**, which are more likely to be functionally important.

**Fig. 3.**
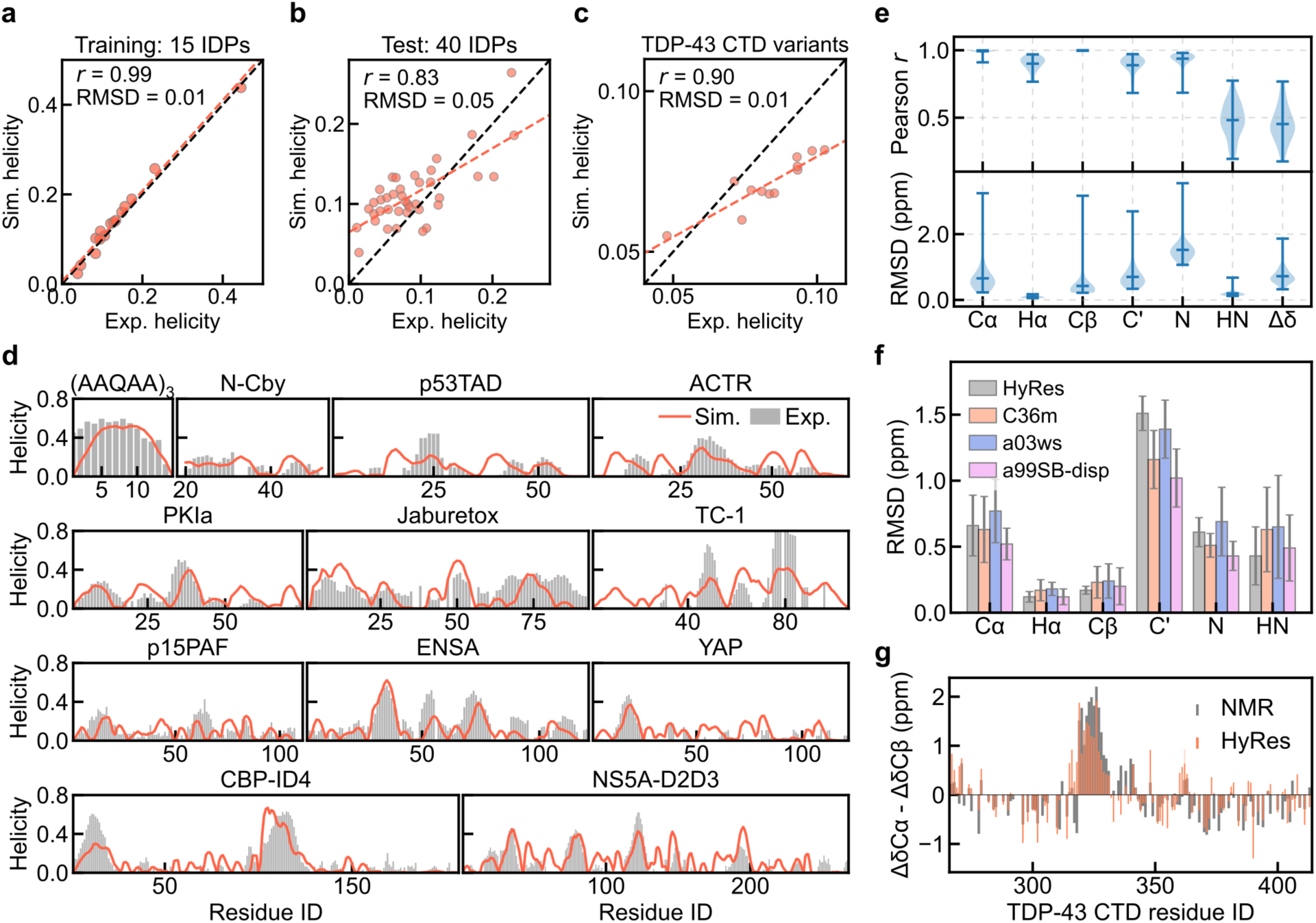
Accuracy of transient local structures in HyRes ensembles. **a-c,** HyRes-predicted mean helicity in comparison to the estimation from NMR secondary chemical shift analysis for the training set (**a**), test set (**b**), and TDP-43 CTD and 11 single mutations (**c**) (see **Supplementary Tables S8-10**). **d,** Representative IDPs showing residual helicity profiles predicted by HyRes simulations (red lines) and estimated from NMR chemical shifts (grey bars). In **a-d**, the experimental helicity was estimated using SSP scores derived from chemical shifts. **e,** Summary of HyRes performance in predicting various chemical shifts and secondary chemical shifts (Δδ = ΔδCα - ΔδCβ), evaluated across 58 IDPs (**Supplementary Table S7**). **f,** Comparison of HyRes with three state-of-the-art all-atom force fields in predicting chemical shifts for ACTR, Ntail, α-synuclein, PaaA2, p15PAF, Sic1, and Ash1 (**Supplementary Table S11**). **g,** Comparison between HyRes predicted and NMR-derived secondary shifts (ΔδCα - ΔδCβ) for TDP-43 CTD.

We also directly examine the ability of HyRes-derived atomistic ensembles to reproduce NMR chemical shifts. Using the TDP-43 CTD as a representative example, HyRes reproduces chemical shifts with high accuracy, yielding correlations of 0.95-1.0 for carbon and nitrogen atoms (**Extended Data Fig. 5b-g**). Similar levels of accuracy were observed across the full dataset of 58 IDPs (**Fig. 3e** and **Supplementary Table S7**), with RMSD below 1 ppm for most atom types. Benchmarked against previously reported results of using all-atom force fields C36m, a03ws, and a99SB-disp^35^ on seven IDPs (**Fig. 3f** and **Supplementary Table S11**), HyRes demonstrates performance comparable to these three state-of-the-art all-atom force fields across all the backbone atoms and Cβ, and slightly better for amide hydrogens (HN).

We further evaluated secondary NMR chemical shifts predicted by HyRes, which report on secondary structure propensities at the residue level besides helicity. Here, secondary shifts were calculated as Δδ = Δδ_Cα_ – Δδ_Cβ_, with Δδ_Cα_ and Δδ_Cβ_ being deviations from the Poulsen IDP/IUP random coil reference values^71^. As illustrated in **Fig. 3g** using TDP-43 CTD, HyRes correctly identifies both positive and negative Δδ in addition to the major helical region (positive Δδ) (**Fig. 3g**), yielding an overall correlation of *r* ∼ 0.77. In-depth analysis of 11 TDP-43 CTD variants demonstrated that HyRes accurately captures single mutation-induced changes in secondary structural propensity (**Extended Data Fig. 6b**). Across the large set of 58 IDPs with available NMR data, HyRes achieves an overall correlation of ∼0.5 and an average RMSD of 0.5 ppm (Δδ in **Fig. 3e** and **Supplementary Table S7**). We note that deviations from experiment mainly arise from residues with negative secondary shifts, suggesting that further refinement could improve the underestimation of extended or β-structures (**Extended Data Fig. 7**), although such states are substantially less populated than transient helices across the analyzed IDPs.

### Dynamic IDP interactions in dilute and concentrated environments

Reliable simulation of dynamic IDP interactions in complex cell-like environments is crucial in studies of IDP function and regulation. The transferability of HyRes to environments beyond the monomeric, dilute condition is first evaluated using the dynamic complex of histone H1 and prothymosin-α (ProTα) in both dilute and condensed phases. These two proteins are heavily charged (+53 for H1 and −44 for ProTα) and have minimal residual structures (**Fig. 4a**). The complex exhibits ultrahigh binding affinity facilitated by extremely fast association kinetics, which enables reversible dissociation on biologically relevant timescales while maintaining tight binding critical for chromatin dynamics^10^.

**Fig. 4.**
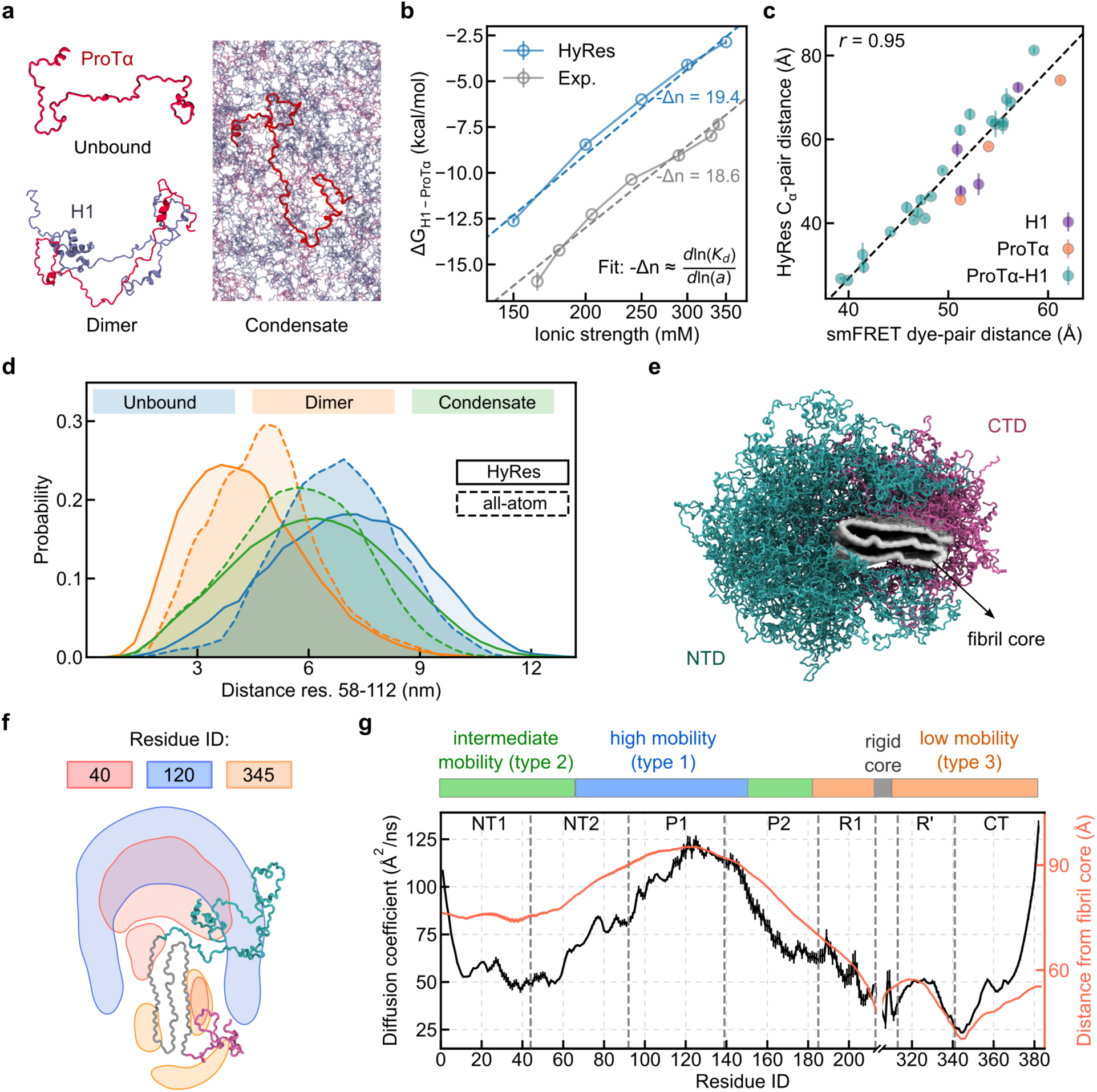
HyRes simulation of dynamic IDP interactions in complex environments. **a,** Representative conformations of unbound ProTα, H1-ProTα dimer, and ProTα within the H1-ProTα condensate. **b,** H1-ProTα binding free energies as a function of ionic strength in the dilute condition calculated using HyRes (blue circles) in comparison to the experimental values (grey circles)^10^. Dashed lines indicate linear fits of - *Δn* = *d*ln(*K*d)/*d*ln(*a*), with *K*d the dissociation constant and *a* the ion activity approximated by ionic strength. **c,** HyRes-derived residue-pair Cα distances vs smFRET-derived dye-pair distances^10^ for the H1-ProTα dimer in the dilute phase. Residue pairs within H1, within ProTα, and between H1 and ProTα are shown in purple, orange, and teal, respectively. In **b** and **c**, error bars for simulation denote block-estimated deviations. **d,** Distributions of ProTα residues 58 and 112 separation in the unbound, dimeric, and condensate states obtained from HyRes in comparison to reported results from all-atom simulations^72^. **e,** A representative structure of the 100-layer amyloid fibril of the full-length hTau in the top view, highlighting the rigid fibril core (grey) and the disordered N- (cyan) and C-terminal (mauve) fuzzy coat regions. **f,** Spatial distributions of residues 40 (red), 120 (blue), and 345 (orange) relative to the fibril core (shown in grey trace). The areas with contour levels of >0.005 were shown. **g,** Residue-specific diffusion coefficients (black) and radial distance from the fibril core (red), illustrating the heterogeneous mobility of the fuzzy coat. Error bars represent deviations estimated from the central 50 monomers of the 100-layer fibril segment. The colored bar above corresponds to assigned residue mobilities from solid-state NMR experiments^73^.

We first performed umbrella sampling to calculate the absolute binding affinity (*ΔG*) of the H1-ProTα dimer in the dilute condition at different ionic strengths (see **Methods** and **Extended Data Fig. 8a-b**). The results show that HyRes captures the nature of extremely tight binding (*ΔG* = - 12.5 kcal/mol at 150 mM salt), but systematically under-estimates experimental values by ∼3 kcal/mol (**Fig. 4b**). This can be mainly attributed to the Debye-Hückel treatment of electrostatic interactions, which under-stabilizes salt-bridge contacts. Nonetheless, HyRes accurately captures the ionic-strength dependence of the binding affinity. This dependence was quantified using the approximated relation of -*Δn* ≈ *d*ln(*K*_d_)/*d*ln(*a*), where –*Δn* corresponds to the number of monovalent counterions released upon association of the two proteins. HyRes yields a value of – *Δn* ≈ 19.4, in close agreement with the experimental estimate of 18.5. We further examined the conformational properties of the H1-ProTα complex in dilute conditions by analyzing residue-pair distances within the dimer and comparing to an extensive set of 30 smFRET measurements on both within H1 and ProTα and across two proteins (**Supplementary Table S12**)^10^. The results reveal a very strong correlation of *r* ∼ 0.95 (**Fig. 4c**), suggesting that the dynamic ensembles of the complex generated by HyRes are highly realistic. The ability of HyRes to accurately capture both free energy and structural details of the H1-ProTα fuzzy complex is remarkable, strongly supporting that HyRes is well-balanced to capture both IDP monomeric conformations and intermolecular interactions under dilute conditions.

To further assess the ability of HyRes to model IDP structure and interaction under concentrated conditions, we examine the conformational ensembles of H1 and ProTα within their condensate containing 80 H1 and 96 ProTα molecules (see **Methods** and **Extended Data Fig. 8c**). The results reveal that both H1 and 96 ProTα exhibit more extended conformations comparing to the complex in the dilute phase, but less extended than unbound ones (**Extended Data Fig. 8d-e**), highly consistent with the experimental observations^72^. Further contact analyses reveal similar intermolecular interactions driving the formation of dimer and condensate (**Extended Data Fig. 8f**). A previous 6-µs all-atom simulation also suggested that the distance between ProTα residues 58 and 112 in the H1/ProTα condensate is smaller than that in the unbound state, but larger than in the dimeric state^72^ (**Fig. 4d**). Note that the 4-µs HyRes simulation (162,256 beads) required only ∼5 days on a single NVIDIA L40s GPU, whereas the all-atom simulations (4,000,932 atoms) required ∼6 months on 36 supercomputer nodes, each equipped with a 12-core CPU and an NVIDIA Tesla P100 GPU. Despite ∼1000-fold higher computational costs, protein conformational distributions remain poorly converged in the all-atom simulation, while those derived from the HyRes simulations are fully converged (**Extended Data Fig. 8g-h**).

### Structure and dynamics of the full-length tau amyloid fibril fuzzy coat

To further assess the ability of HyRes in modeling dynamic proteins in complex environments, we investigate the “fuzzy coat” of the amyloid fibril of the full-length protein tau^74^ (**Fig. 4e**). Fuzzy coats, formed by disordered segments not part of the structured fibril cores, are the primary interface for interactions with cellular partners and have been recognized as the central regulator of fibril toxicity, seeding activity, and functional modulation^75^. Studies of the fuzzy coats are extremely challenging for atomistic simulations both due to the system size and the complex crowded environment^76,77^, where achieving convergence is rarely feasible. We constructed a 50 nm-long fibril segment containing 100 layers of the full-length 383-residue phospho-mimetic human tau (hTau; 0N4R) protein (see **Methods** and **Extended Data Fig. 9a**). Note that only residues 215-304 form the structured core (PDB: 8TTN), and the N- and C-terminal disordered domains (NTD and CTD) are 214 and 79 residues long, respectively. Nonetheless, good convergence was achieved for both the dimension and secondary structures of each monomer with only a 4-µs HyRes simulation, which required ∼5 days on a single L40S GPU (**Extended Data Fig. 9b-c**). This is extraordinary, highlighting the efficacy of HyRes for modeling dynamic proteins in the ultra-crowded local environment surrounding the fibril core.

Further analysis reveals that NTD exhibits a smaller Flory polymer scaling exponent than the short C-terminal domain (CTD) and narrower distributions for both *R*_g_ and *R*_ee_. (**Extended Data Fig. 9e-g**). Curiously, NTD has a smaller average *R*_ee_ although being ∼3-fold longer than CTD. Analysis of the spatial distributions of individual residues relative to the fibril core reveals the overall architecture of the fuzzy coat. It shows that NTD adopts loop-like conformations, where the NT1 and part of the NT2 regions fold back toward the fibril core surface, thereby exposing the P1 domain as the outermost layer of the fibril (**Fig. 4f-g**). In contrast, the C-terminal domain (R’ and CT regions) resides closer to the fibril core, and exhibits stronger interactions with the core (**Extended Data Fig. 9d**). As a result, there is a strong correlation between residue-wise diffusion coefficients and spatial positioning (black vs red curves in **Fig. 4g**). These dynamic features are highly consistent with a recent solid-state NMR study^73^, which suggested three distinct mobility regimes (top color bar in **Fig. 4g**). The ability of HyRes simulations to recapitulate heterogeneous dynamic properties across the fuzzy coat further supports the robustness of HyRes in capturing structure and dynamics of IDPs in highly crowded and structurally complex environments.

### Proteome-scale high-resolution disordered conformational ensembles

Leveraging the efficiency and accuracy of HyRes, we applied it to generate high-resolution disordered conformational ensembles for all 29,872 IDPs (including intrinsically disordered regions or IDRs) from the human IDRome and DisProt databases (see **Methods**). The proteome-scale simulations allow the creation of an open-access HyRes-IDRome database containing all resulting HyRes conformational ensembles, as well as key conformational and sequential features such as *R*_g_, R_ee_, average residual helicity, helical segment information, and Flory scaling exponent (see **Methods**). With extensive validations as described above, the HyRes-IDRome database can serve as a robust resource for further developing better machine learning models and for deeply understanding the sequence-ensemble-function-disease relationships. Note that all-atom side chains can be readily reconstructed as described above.

Compared to CALVODOS-derived IDRome database^30^, we observe high correlations (*r* = 0.99) for both *R*_g_ and *R*_ee_ (**Extended Data Fig. 10a-b**), which is expected given comparable abilities of these two models to reproduce *R*_g_ of known IDPs (**Fig. 1e**). Most IDPs behave as random coils with a mean scaling exponent of 0.54, while significant fractions exhibit highly compacted (*v* < 0.4) or highly expanded states (*v* > 0.7) (**Fig. 5a**). There is a wide distribution of helical propensities, with an average α-helicity of ∼0.1 and a notable fraction of sequences exhibiting propensities exceeding 0.2 (**Fig. 5b**). Further analysis of transient helical segments resolves a broad distribution of helical segments exceeding 25 residues in length and up to 80 residues (**Fig. 5c** and **Extended Data Fig. 10e-f**). There is a positive correlation between the average populations of the helical segments and their lengths (**Extended Data Fig. 10c**). Furthermore, analysis of amino acid composition across all IDP sequences and within the identified helical segments reveals that the helix-disrupting residues glycine, proline, and serine are depleted in helices, while alanine, glutamic acid, and leucine are enriched (**Extended Data Fig. 10d**). No significant enrichment or depletion was observed for the remaining amino acids.

**Fig. 5.**
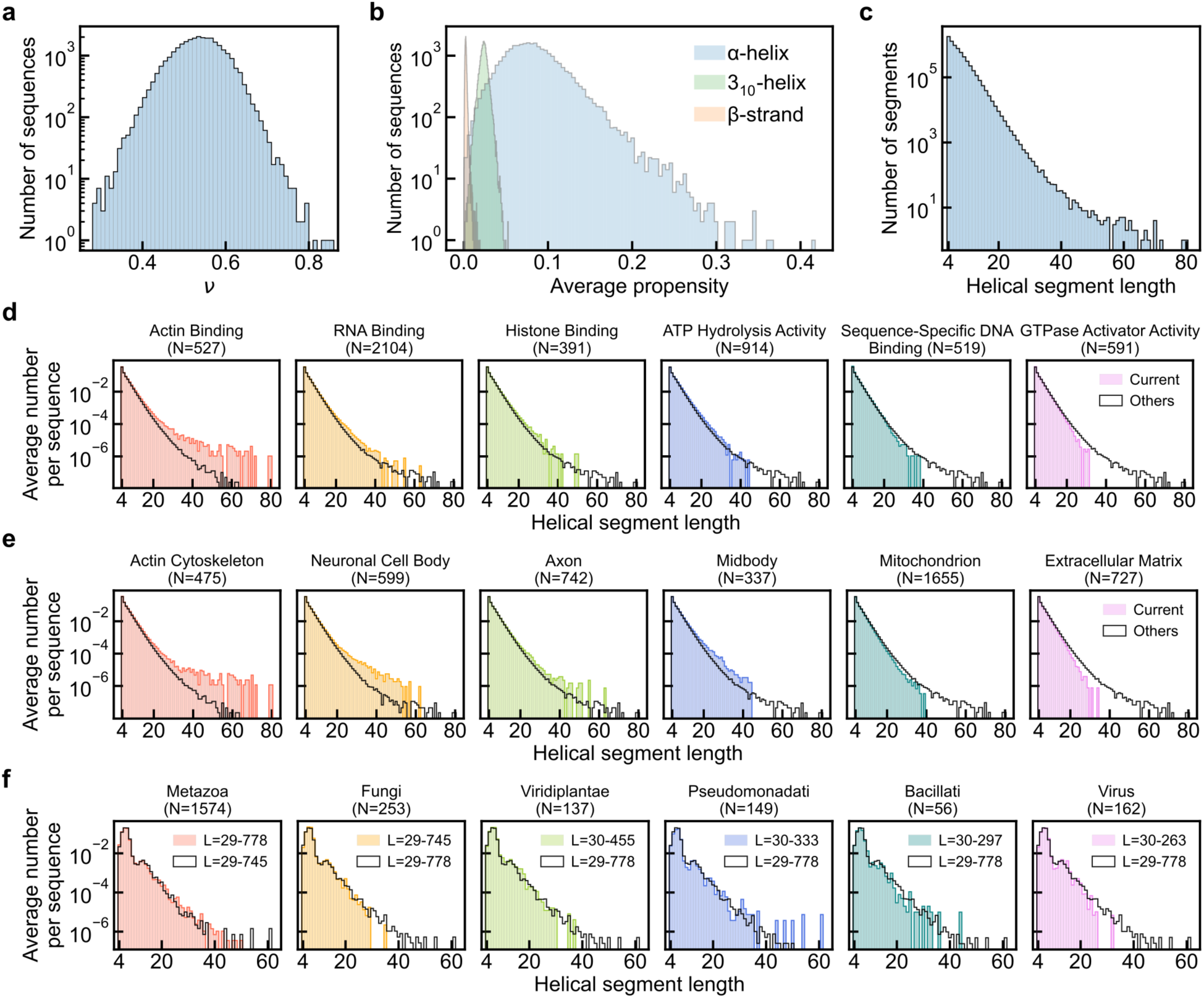
Proteome-scale HyRes simulation of IDPs. **a**, Distribution of the Flory scaling exponent, *ν*, for all 29,872 IDPs from the human IDRome and DisProt databases. **b**, Distributions of average secondary structural propensities. **c**, Distribution of helical segment length. **d-f**, Average number of helical segments per sequence for proteins associated with various functions (**d**), cellular locations (**e**), and organism groups (**f**). The number of sequences (N) and the sequence length range (L) are indicated. The unfilled black step plots are the distributions of all the remaining sequences. The entire sequences are from IDRome with length in the range of 30-999 for panels **d** and **e**, and from DisProt for panel **f**.

We further compared the local helical segment appearance among various functional, location, and taxonomic classifications (**Fig. 5d-f**) using the average number of helical segments per sequence (HS_avg_). For shorter helices (< 20 residues), a strikingly similar level of HS_avg_ is found across different molecular functions, cellular locations, and organism groups. However, substantial diversity can be observed for the presence of longer helices among human IDPs belonging to different functional groups or associated with different cellular locations. For instance, actin- and RNA-binding IDPs, as well as those localized to the actin cytoskeleton, neuronal cell body, and the axon, exhibit a markedly higher propensity to form extended local helices. Conversely, sequences involved in GTPase activator activity or associated with the extracellular matrix show a sharp depletion of these longer features. Interestingly, the distribution of helical segments is remarkably similar across various organism groups (**Fig. 5f**), with only minor differences observed for the appearance of helical segments longer than 30 residues.

## Discussion

In this work, we have introduced HyRes, a well-balanced, physics-based, hybrid-resolution protein force field that resolves a long-standing bottleneck in the structural biology of IDPs. HyRes features an explicit atomistic backbone with intermediate-resolution sidechains, and employs a physics-based energy function that includes bonded terms, van der Waals interactions, electrostatics, and backbone-mediated hydrogen bonds. It bridges the gap between atomic accuracy and extreme computational efficiency. Through extensive cross-validation against a benchmark of approximately 100 IDPs, we demonstrate that HyRes generates atomistic ensembles that match or outperform state-of-the-art all-atom force fields in reproducing experimental chain dimensions, transient long-range contacts, and local secondary structures, but at several orders of magnitude lower computational cost.

Leveraging this unique combination of accuracy and scalability, we have generated detailed conformational ensembles for nearly 30,000 IDPs across the human proteome and the DisProt database. A major foundational strength of this massive resource is that it represents the first proteome-scale collection of atomistic disordered ensembles generated entirely via a physics-based approach that has been rigorously validated against virtually all available experimental IDP data. Our initial analysis of this database already reveals striking, sequence-encoded correlations between residual structural properties and specific cellular functions or localizations, providing a fresh paradigm for understanding how IDPs fine-tune cellular regulation.

A critical distinction of the HyRes physics-based molecular simulation framework is its outstanding environmental transferability. Unlike machine learning models that are strictly constrained by their training conditions, HyRes is grounded in fundamental physical forces. This enables the same underlying model to accurately capture dynamic, sequence-dependent IDP behavior across vastly different biological environments, ranging from isolated monomers in dilute phases to dense, multivalent networks in biomolecular condensates and the crowded “fuzzy coats” of amyloid fibrils. Ultimately, HyRes and the comprehensive, open-access HyRes-IDRome database provide an unprecedented structural resource for the molecular biology community. By offering an efficient, physics-governed window into the disordered proteome, this work not only opens new avenues for studying IDP function and disease pathogenesis but also establishes a rich, high-fidelity structural foundation to accelerate deep learning applications in protein science.

## Methods

### HyRes optimization through hierarchical rebalancing of interactions

The HyRes model features an atomistic backbone and intermediate CG side chains. The total potential energy contains contributions from bonded, nonbonded, and backbone hydrogen-bond interactions^51–53^. To rebalance the electrostatic and vdW interactions, the relative dielectric constant was determined as *ε*_r_ = 60.0 at room temperature based on the radii of gyration of 30 random and one alternating polypeptides of glutamic acid and lysine, EK25^57^ and (EK) ^58^ (**Extended Data Fig. 1a and b**). With *ε*_r_ = 60.0, *R*_g_ of (EK)_16_ under various ionic strengths can be well captured. Previous benchmarks have shown that ABSINTH overestimates the *R*_g_ by ∼30%^58^. Indeed, scaling down *R*_g_ values derived from ABSINTH by 75% reproduced those predicted by HyRes with *ε*_r_ = 60.0.

The self-interactions between arginine side chains were specifically modulated to reproduce the *R*_g_ and SAXS profile of deca-arginine (R_10_)^59^ (**Extended Data Fig. 1c and d**). Then the vdW interactions for other amino acids were refined using *R*_g_ of 20 A1-LCD variants^60^, during which the relative strengths of cation-π and π-π interactions among positively charged and aromatic amino acids were roughly maintained^78^. Interaction parameters were then iteratively fine-tuned until the best agreement was reached.

To fine-tune the residual helicity, the hydrogen bond interaction strength was first tuned to reproduce the helicity of (AAQAA)_3_. After that, 15 IDPs that have significant transient helical domains were selected to fine-tune the helical propensities of each amino acid type (**Supplementary Table S8**), through modulating special 1-4 interaction radii between side chains and backbone atoms. Due to the relatively mild residual helicity of many residue types, we found it ineffective to maximize the correlation between simulated and NMR-derived values. Instead, we minimized the difference between average residual helicities from simulation and NMR for each amino acid type (**Extended Data Fig. 5a**) as well as the difference between the simulated and NMR-derived average overall helicity of the 15 IDPs in the training set (**Fig. 3d**).

### Curation of various validation sets

These rebalanced parameters were finally validated using the set of 98 IDPs (**Supplementary Table S3**) collected from previously published papers^49,61^ and small angle scattering biological data bank (SASBDB)^79^ filtered using the tag “IDP”. In addition, the measurements under special conditions, including additional osmolytes (*e.g.*, urea, GdmCl, TMAO), low pH (< 6.5), and multivalent ions (like Zn^2+^ and polyamines), were also excluded. For comparing scattering profiles, we employed the subset containing 45 IDPs with high-quality SAXS data, which showed minimal hints of aggregation and were previously re-analyzed and validated using the molecular form factor approach^49^. For the validation of chemical shifts and secondary chemical shifts, entries in DisProt^4^ were filtered using the tag “NMR”. The items stored BMRB cross-references were selected. Measurements involving special conditions (*e.g.*, see above) or including folded domains were excluded. At the end, we collected 58 IDPs as summarized in **Supplementary Table S7**. Additional helicity profiles were collected from published works for those without NMR raw data (sequences without BMRB IDs in **Supplementary Tables S8-9**).

### HyRes simulation and analysis of monomeric IDPs

All HyRes simulations were performed on GPUs using OpenMM^80^ and HyresBuilder (https://github.com/lslumass/HyresBuilder/) packages. Langevin dynamics was performed with a collision frequency of 0.1 ps^-1^ and a time step of 8 fs. All bonds involving hydrogen atoms were constrained by the SHAKE algorithm. The cut-off method was applied for nonbonded interactions with a cutoff of 1.2 nm for Lenard-Jones interactions and 1.8 nm for Debye-Hückel electrostatic interactions, which were smoothly switched off from 1.0 and 1.6 nm, respectively.

For single-chain simulations, a linear chain was constructed using HyresBuilder, followed by a quick minimization and equilibration. Under the consistent temperature and ionic strength of experimental conditions, at least 2 µs of production simulation was performed to calculate the properties of each IDP. MDAnalysis^81,82^ was used to calculate the radius of gyration, end-to-end distance, secondary structure propensity (implemented DSSP^70^), and any other analysis related to the trajectory. GROMACS “gmx saxs”^83^ was used to estimate the scattering profiles. For calculating NMR-related properties, HyRes simulation trajectories were first back-mapped into atomistic ones using cg2all^67^ with the backbone model. Using these atomistic trajectories, chemical shifts were estimated by SHIFTX2^68^. To calculate secondary shifts, Poulsen IDP/IUP random coil chemical shifts were predicted as the references^71^. PRE profiles were estimated using DEER-PREdict^65^ from each atomistic ensemble, during which the tumbling time *τ*_c_ was optimized in the range of 1-7 ns (*τ*_t_ = 1 ps) to obtain minimum deviation from experimental data. Three replicas were performed for a better estimation of PRE profiles.

The Flory scaling exponent, *ν*, was estimated based on the intramolecular structure factor, *ω*(*q*), calculating through^84^

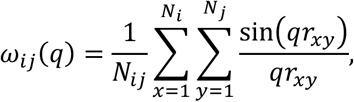

where *N_i_* is the number of beads of type *i*. Cα is selected for the calculation; thus, *N_ij_* = *N_i_* (*i* = *j*). *ν* is related to the slope (= −1/*ν*) of *ω*(*q*) over the intermediate scaling regime, which can be estimated as 2π/*R*_ee_ << *q* << 2π/*b*, where *b* is the Kuhn segment. To quantify the helical segments found from IDP ensembles, the helical segments longer than 4 residues were counted frame by frame for each ensemble. For any two segments, if their starting and ending residues are similar (residue ID difference is smaller than 4), they are recognized as the same helical segment. All these segments are counted together as one segment using their average length.

### Fuzzy complex of H1-ProTα: simulation and free energy analysis

ProTα monomer was constructed as a random coil using HyresBuilder. The H1 folded domain was converted from the structure of PDB 6HQ1^85^. During the simulations, this folded domain was treated as a rigid body to maintain the structure. For the H1- ProTα dimer simulation, monomers were randomly placed in the simulation box, and a 4 µs-long simulation was performed to sample the dimer state. A random dimer state was selected to initiate umbrella sampling to estimate the binding free energy under various ionic strengths. For each umbrella sampling simulation, reaction coordinates were defined as the distance between the centers of mass of H1 and ProTα, and 50 sampling windows were performed with minima at 0.5 to 25.5 nm in steps of 0.5 nm. Each replica was performed for 2 µs with the force constant of 100 kJ/mol/nm^-2^. Weighted histogram analysis^86^ was used to determine the potential of mean force (PMF) using the program wham^87^. For the H1-ProTα condensate simulation, the pre-compacted condensate was first prepared through NPT simulation, resulting in a 27-nm cubic box containing 96 ProTα and 80 H1 molecules. The slab configuration was subsequently built by extending the box edge to 45 nm in the x direction, and then relaxed, and a 4 µs-long HyRes simulation was performed. The distribution of the residue 58-112 distance was collected from all ProTα molecules.

### Amyloid fibril of full-length tau: fuzzy coat modeling and simulation

Starting from the structure of the tau fibril core (PDB 8TTN), the core region was extended to 100 layers to build a 50 nm-long fibril segment. The missing disordered regions were then built using Modeller^88^, followed by a quick relaxation (**Supplementary Fig. S1a**). The final system contains 231,300 particles, equivalent to 563,800 atoms without water in all-atom simulations. In the HyRes simulation of the fibril of the full-length tau, the core domains were maintained using positional restraints, while the rest of the tau proteins were free. The production simulation was performed for 4 µs to equilibrate and sample the dynamic ensembles of the fuzzy coat. Due to the highly crowded fuzzy coat environment, the effective dielectric constant within this region is expected to be lower than that of a dilute solution, even though the precise value has not been determined. As such, we simulated the full-length tau fibril using the dielectric constants of 60.0, 40.0, and 20.0, indicating that the overall feature of fuzzy coat arrangement and local diffusion patterns were identical (**Supplementary Fig. 1b**). To minimize the finite-size effect, only the 50 monomers in the central region of the fibril were selected for analysis.

### HyRes-IDRome database

#### Sequence collection

The sequences used to construct the open-access database were selected from DisProt (release 2025_06 with ambiguous evidence) and the human IDRome database^30^. DisProt contains 13,347 entries, from which duplicate sequences were merged and only one representative was retained. Afterwards, the IUPred2A^89^ tool was used to get the disorder prediction score, and sequences were retained if the residue score was larger than 0.5. In addition, sequences containing three or fewer ordered residues were also included. Furthermore, sequences shorter than 30 residues or longer than 1000 residues and sequences with any non-canonical residues were discarded. Finally, we collected 2,383 sequences from DisProt, corresponding to 1,218 unique proteins, of which 557 are unique human proteins. Following the same protocol, 27,833 sequences were selected from IDRome. We further compared them to the newly released UniProt (release 2026_01) and removed the items that were deleted to finally collect 27,489 sequences, corresponding to 15,179 unique proteins. Together, the final HyRes-IDRome database contains a total of 29,872 unique IDP sequences.

#### Database configuration

HyRes simulations were performed for 2 µs and 4 µs for the sequences shorter and longer than 300 residues, respectively. To assess the convergence of the HyRes simulation, four sequences of varying length were selected randomly. For each sequence, the simulation trajectory was divided into equal halves, and both radius of gyration and helicity were calculated for each half. Similar distributions of *R*_g_ and helicity obtained from the first and second halves indicate the simulations are converged for various lengths (**Supplementary Fig. S2**). For each sequence, 1,000 frames were extracted from the latter 90% of the trajectories at equal spacing and deposited into the database together with corresponding properties. The latter includes radius of gyration, end-to-end distance, residual secondary structure propensities, and helical segment counting, as well as sequence features derived from localCIDER^90^, including fraction of charged residues, net charge per residue, number of negatively charged residues, number of positively charged residues, number of neutral residues, charge segregation parameter, mean hydropathy, and fraction of each amino acid type. All curated datasets are publicly available through Hugging Face at https://huggingface.co/datasets/mdlab-um/HyRes-IDRome. Users can download individual sequence folders or the entire dataset from the online repository using Git or the Hugging Face Hub Python library. For each sequence, a folder is created which includes a PSF file, a DCD trajectory file, three NumPy files containing the radius of gyration, secondary structures assignments, and end-to-end distances. In addition, an Excel workbook is also provided for each sequence and contains three worksheets, containing sequence-level properties and information, simulation-derived observables, biological annotations, all helical segments, and per-residue average helicity values.

### Taxonomic Information, Molecular Function, and Cellular Location

Taxonomic information for each DisProt sequence was obtained using the NCBI Taxonomy database. The taxon ID provided for each sequence was used to extract taxonomic classifications with the ETE3 tool^91^. At certain taxonomic levels, a small number of sequences were labelled as “Unknown,” indicating that they could not be assigned to a specific category at that level. All sequences labelled as “Unknown” at the domain level were identified as viruses and were therefore classified under the domain *Virus*. Gene Ontology (GO) terms for IDRome sequences were obtained from the GO analysis reported in a previous study conducted by Tesei and colleagues^30^. In total, 68 distinct molecular functions and 94 distinct cellular locations were identified.

## Acknowledgements

The authors acknowledge helpful discussions from Drs. Xiaorong Liu, Yumeng Zhang, and Xiping Gong in the development and optimization of HyRes. This work is supported by NIH through R35 GM144045 (to Chen).

## Author contributions

Li and Chen conceptualized the idea. Li conducted HyRes refinement and benchmarks. Barethiya designed and performed simulation and analysis of the HyRes-IDRome database. Li, Barethiya, and Chen drafted and revised the manuscript.

## Additional information

Extended data is available for this paper at https://doi.org/xxxx

## Supplementary information

The online version contains supplementary material available at https://doi.org/xxxx

Correspondence and requests for materials should be addressed to Jianhan Chen

---

**Extended Data Fig. 1.**
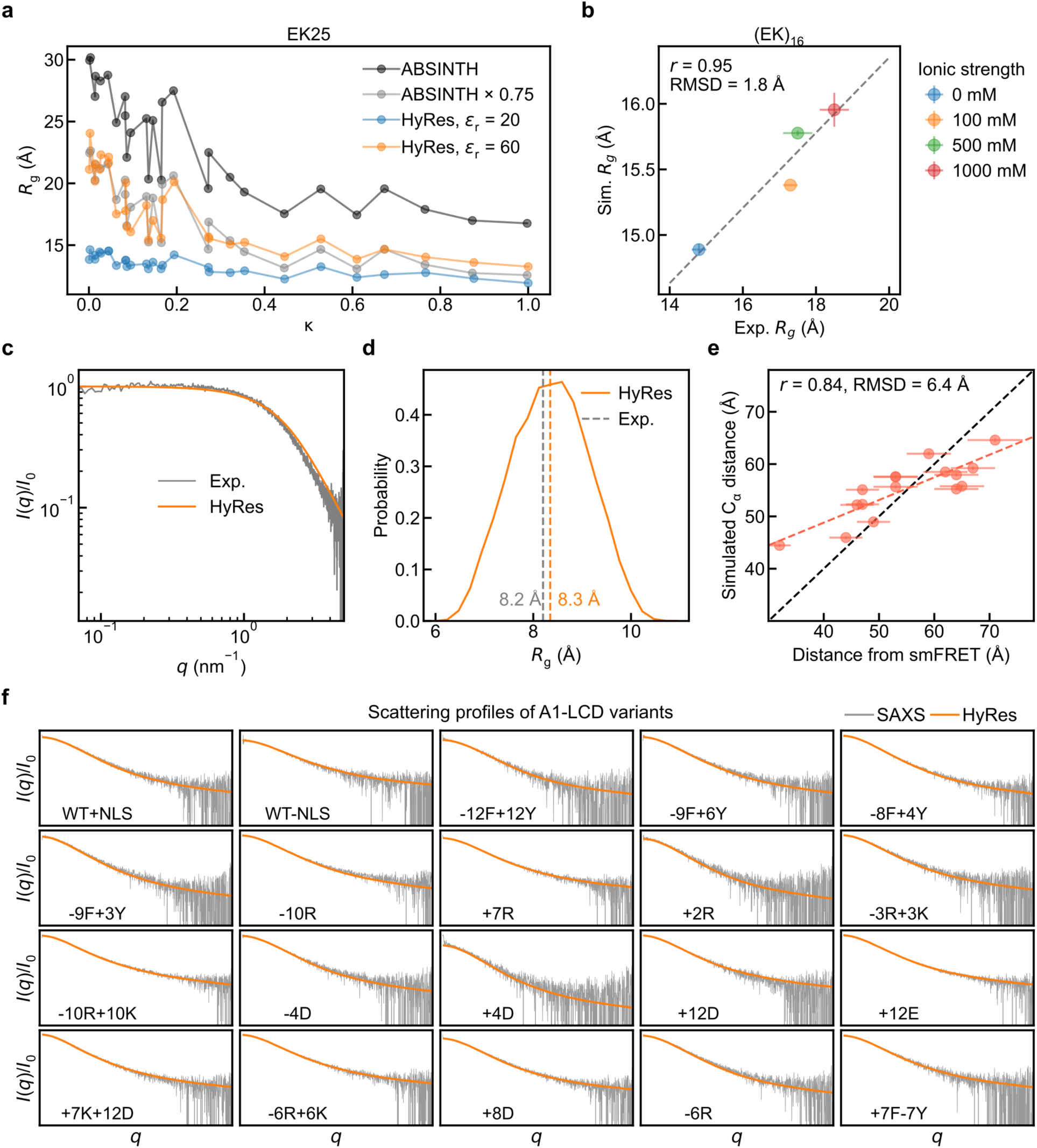
Refinement of the HyRes model. **a**, Comparison of *R*_g_ values of EK25 peptides between HyRes and ABSINTH estimations. **b**, HyRes-derived and SAXS-derived *R*_g_ values of (EK)_16_ under various ionic strengths. **c** and **d**, Comparison of scattering profiles and *R*_g_ values of R10 between HyRes simulation and experiment. **e**, Comparison of residue-pair distances between HyRes simulations and smFRET using the dye pair of Cy3D-CF660R for 16 IDPs as shown in Fig. 1d. **f**, Scattering profiles calculated from HyRes ensembles in comparison with SAXS measurements for 20 A1-LCD variants. Error bars of the simulation-derived values were calculated from the block deviations.

**Extended Data Fig. 2.**
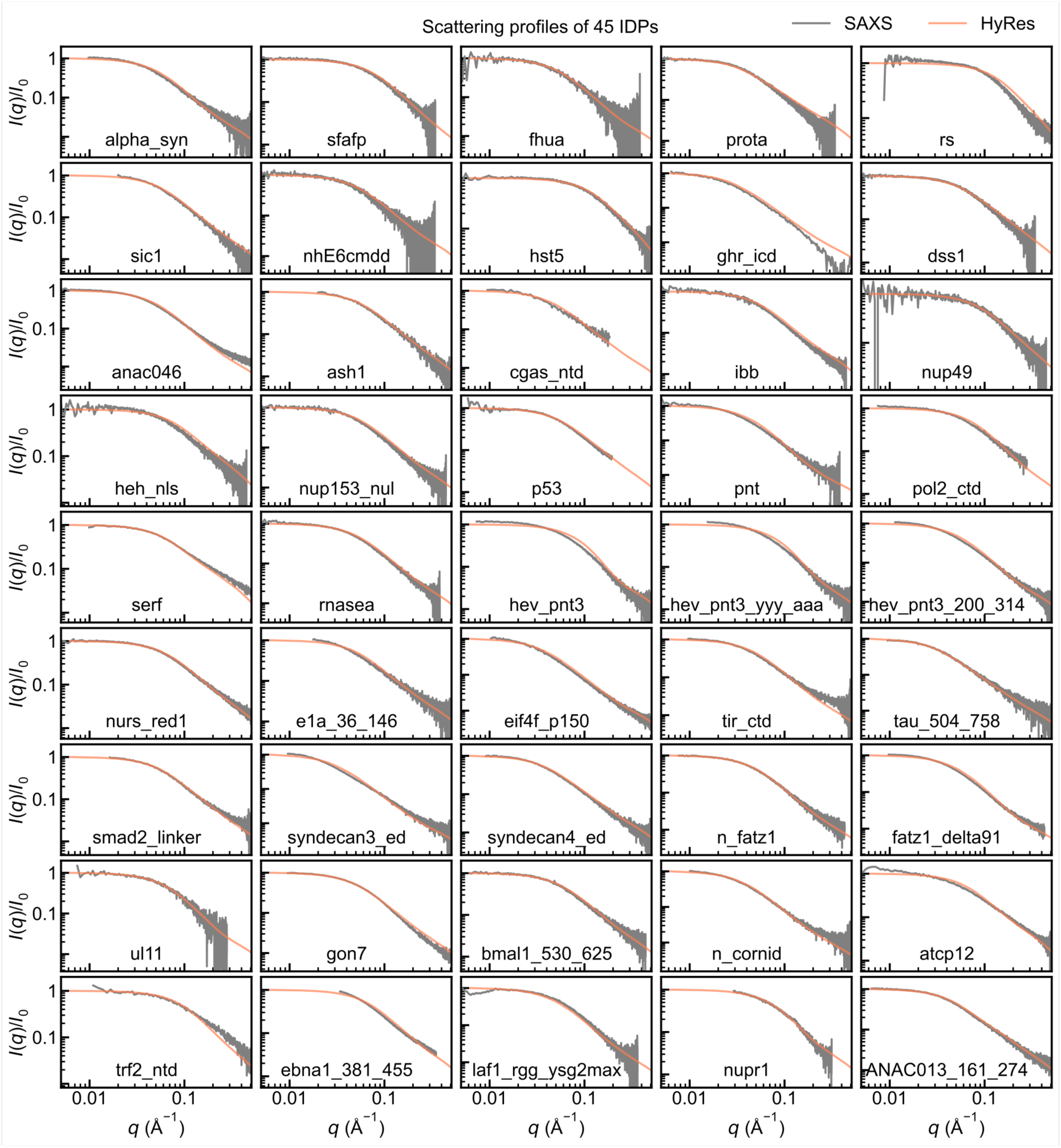
HyRes-derived scattering profiles compared with SAXS data for 45 IDPs.

**Extended Data Fig. 3.**
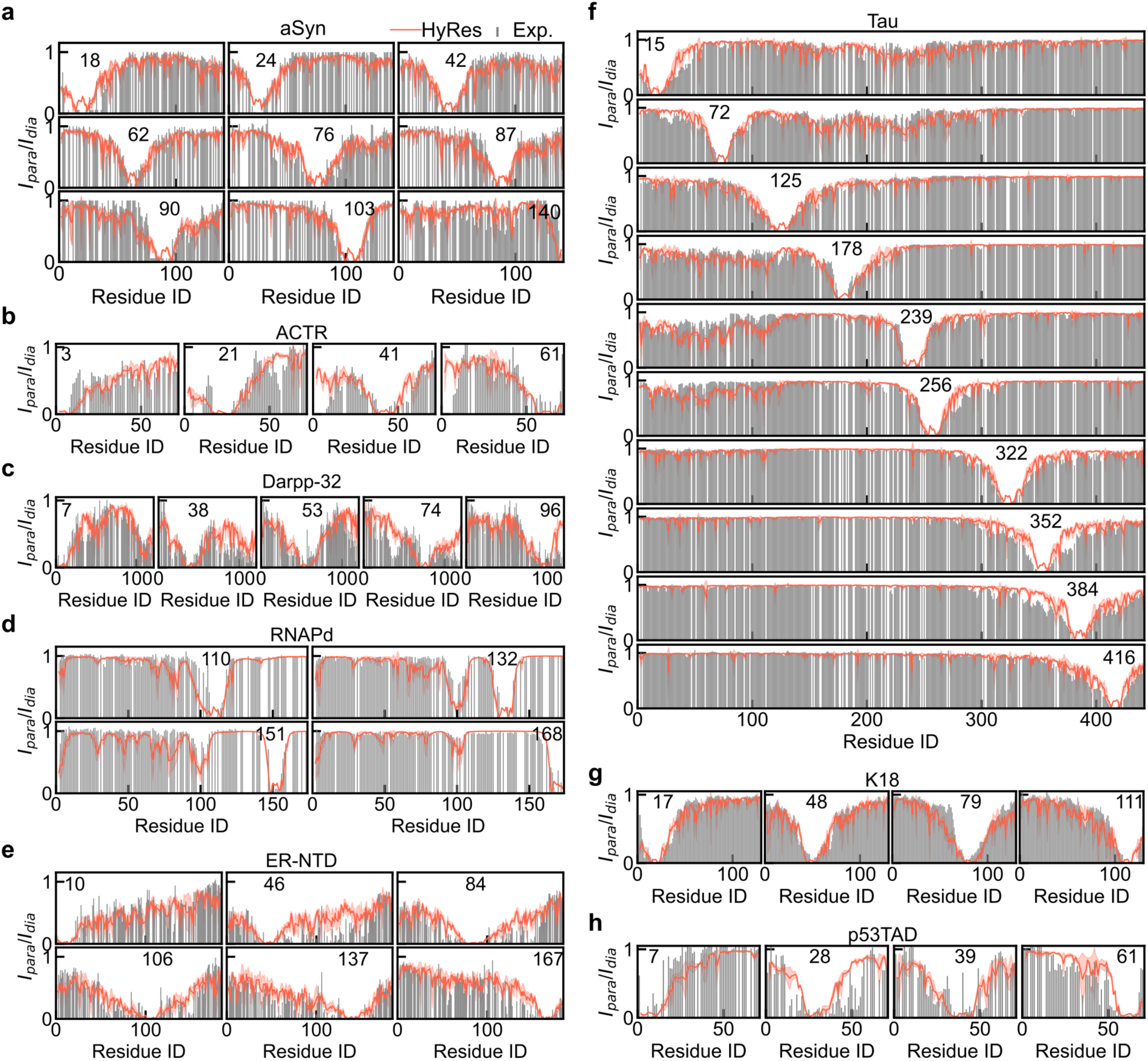
HyRes-derived PRE profiles compared with NMR-derived data for aSyn (**a**), ACTR (**b**), Darpp-32 (**c**), RNAPd (**d**), ER-NTD (**e**), Tau (**f**), K18 (**g**), and p53TAD (**h**). Spin-labeled residues are indicated by residue numbers. Shaded regions represent deviations estimated from three independent replicas.

**Extended Data Fig. 4.**
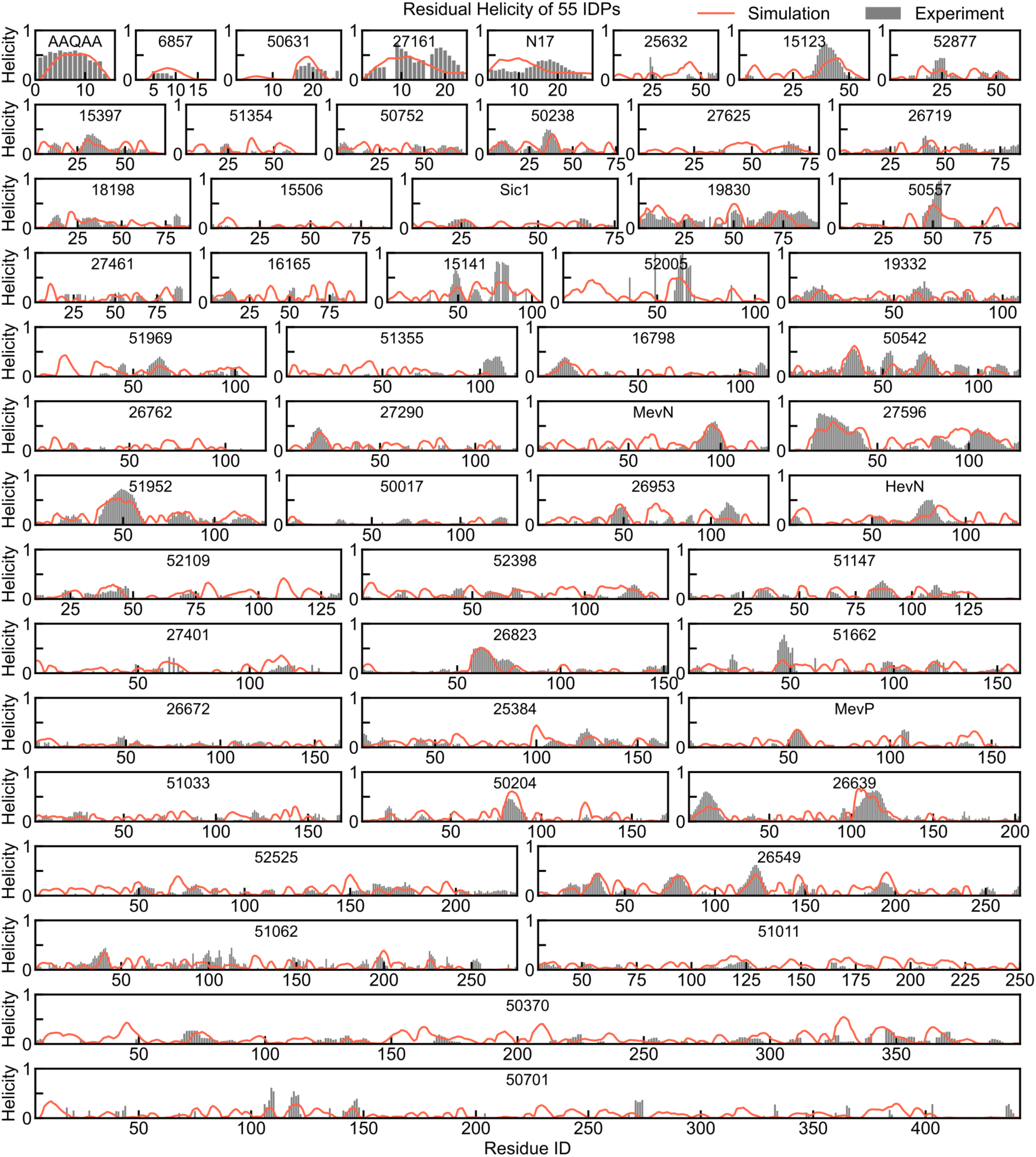
HyRes-derived residual helicities (red lines) in comparison with NMR-derived values (grey bars) for 55 selected IDPs. Corresponding BMRB entry IDs or IDP names are noted.

**Extended Data Fig. 5.**
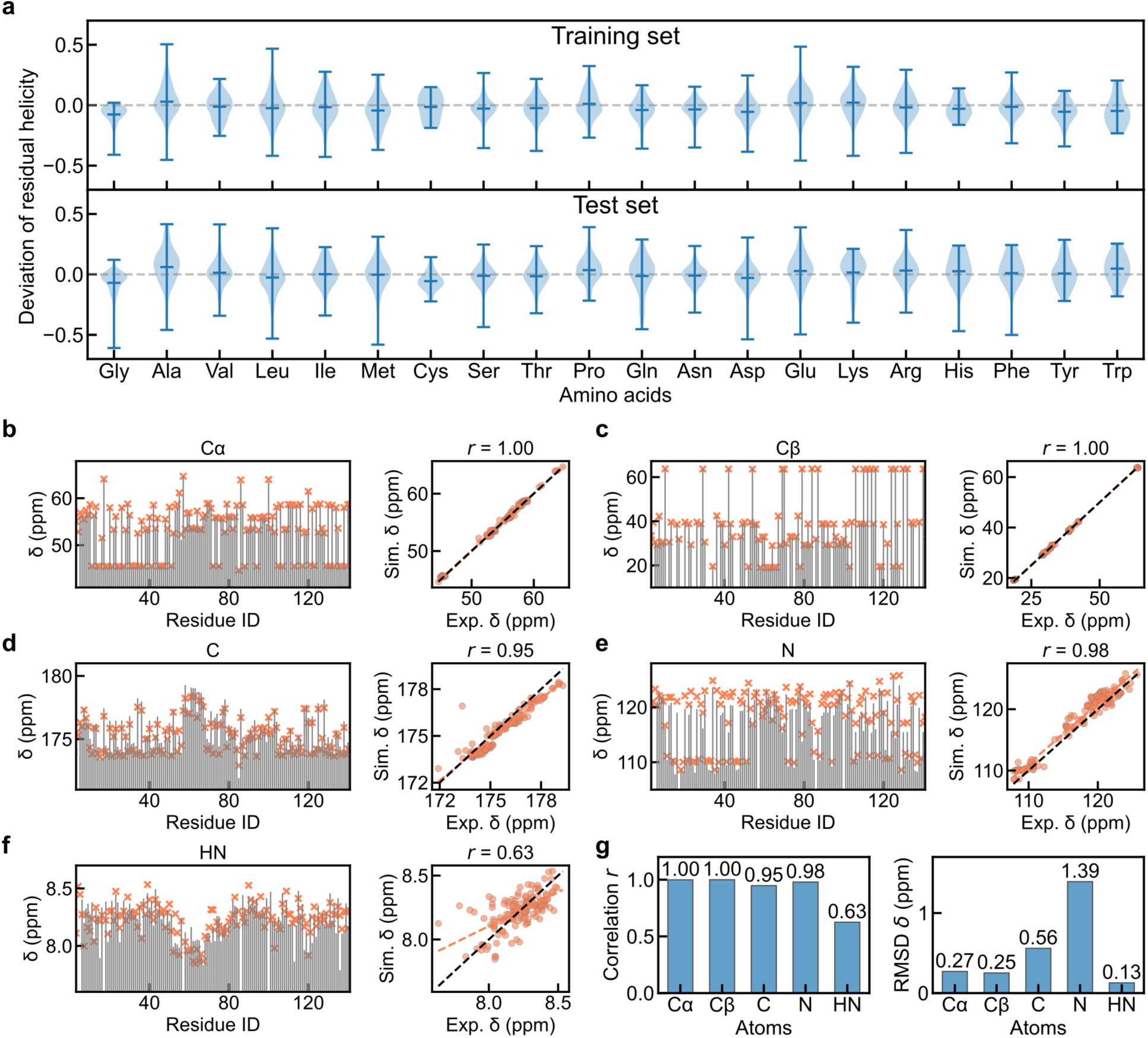
Training and validation of HyRes using residual helicity and NMR chemical shifts, respectively. **a,** Deviations of residual helicities from the estimations from NMR measurements for training and test sets. **b-f**, Comparison of chemical shifts of Cα, Cβ, amide C, N, and HN between HyRes simulations and NMR measurements for TDP-43 CTD. **g**, Correlation and RMSD between HyRes-predicted and measured NMR chemical shifts for TDP-43 CTD.

**Extended Data Fig. 6.**
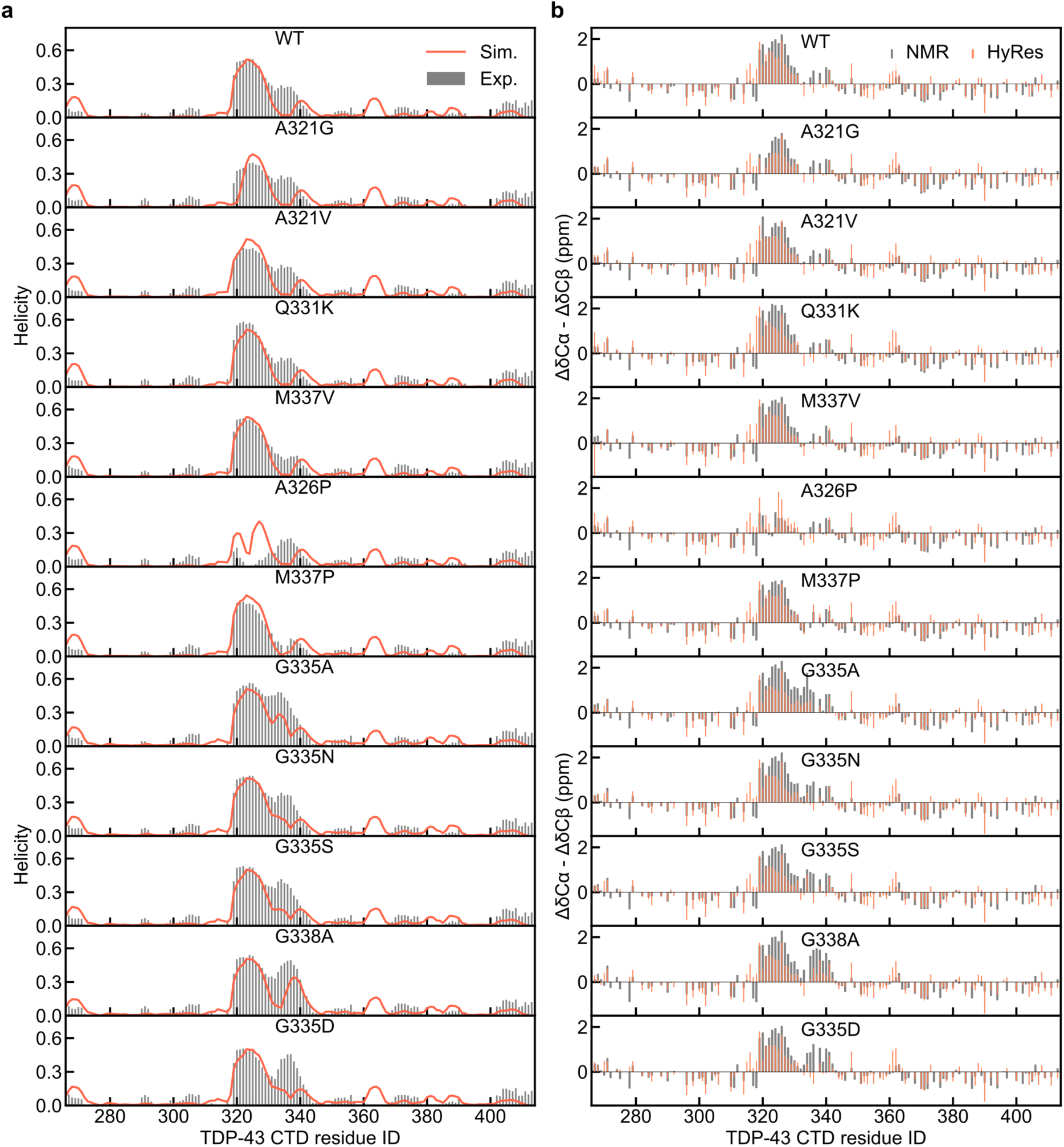
Ability of HyRes to capture the effect of single mutations of TDP-43 CTD for (**a**) residual helicities and (**b**) secondary chemical shifts. HyRes results are shown in red and NMR results in grey bars.

**Extended Data Fig. 7.**
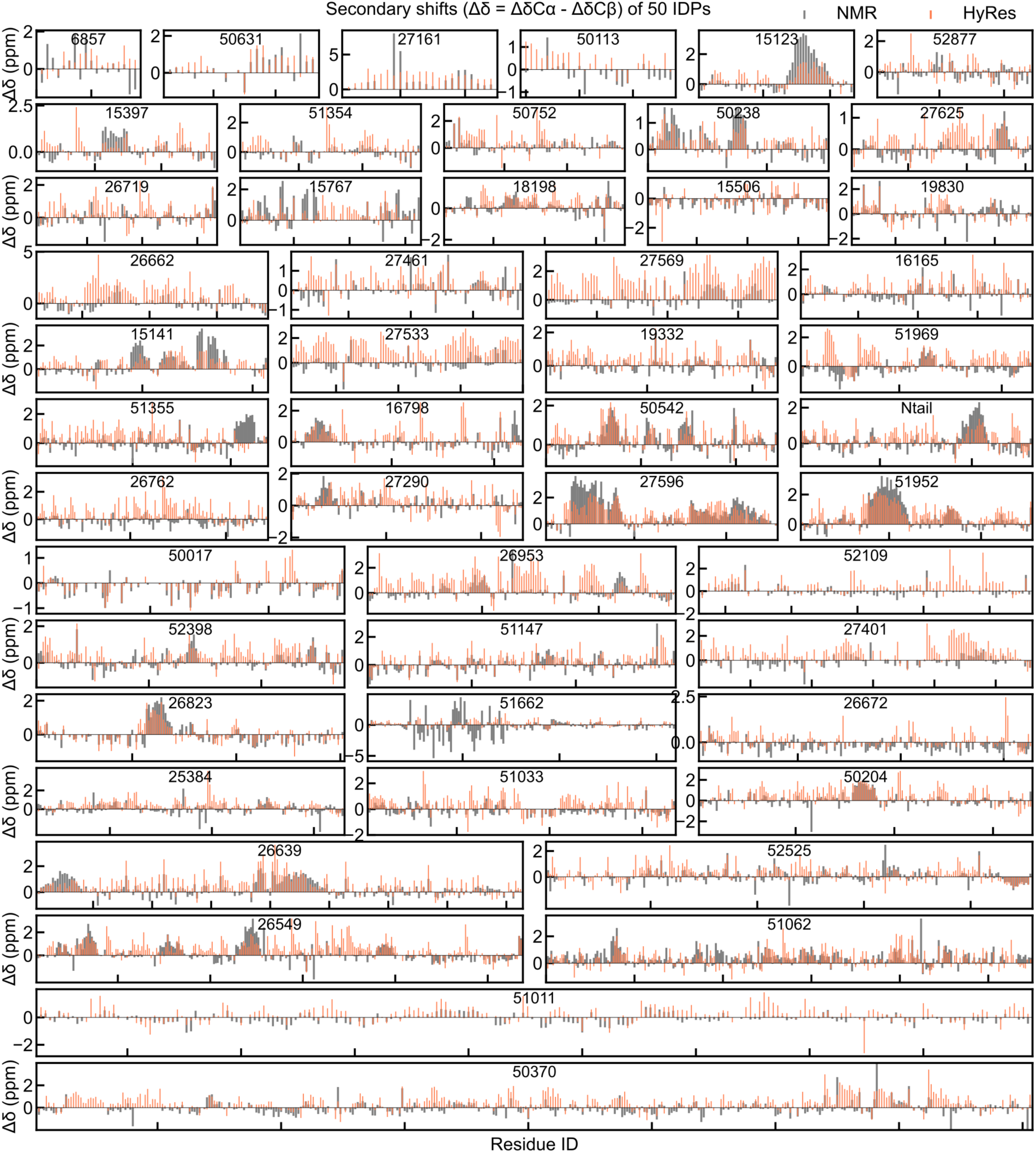
HyRes-derived secondary shifts (red bars) in comparison with NMR-derived values (grey bars) for 50 IDPs (not including TDP-43 variants). Corresponding BMRB entry IDs are noted.

**Extended Data Fig. 8.**
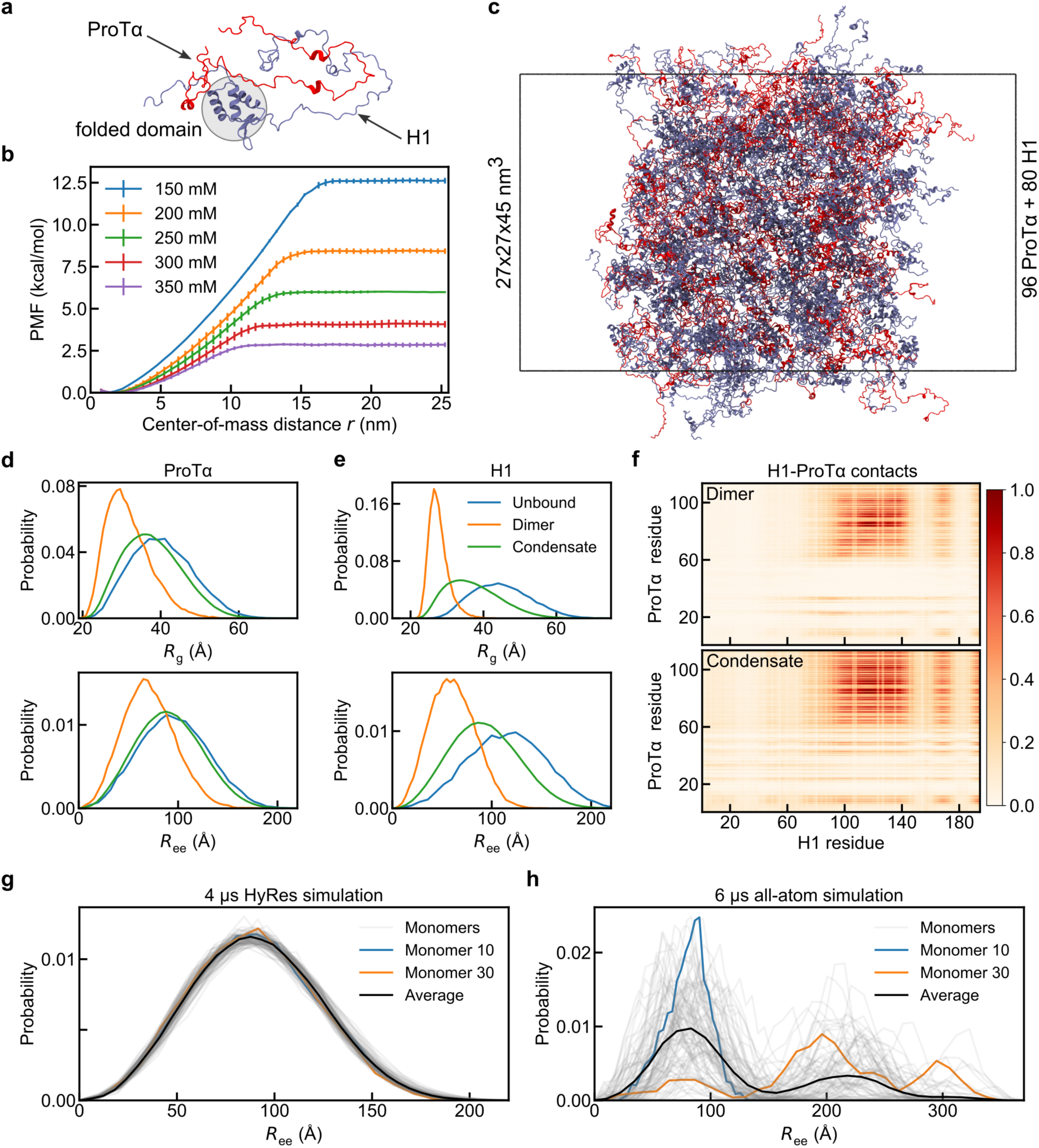
HyRes simulation of H1-ProTα dimer and condensate. **a**, The initial configuration of the H1-ProTα dimer for calculating binding free energy. **b**, PMF profiles of H1-ProTα interaction under increasing ionic strengths calculated from umbrella sampling simulations. **c**, A representative snapshot of the condensate containing 80 H1 (ice blue) and 96 ProTα (red) molecules in a slab configuration. **d**-**e**, Distributions of *R*_g_ and *R*_ee_ of ProTα (**d**) and H1 (**e**) in the unbound, dimer, and condensate states, respectively. **f**, Normalized H1-ProTα contact maps in the dimer and condensate states. Contact densities were normalized based on the maximum value. **g-h**, *R*_ee_ distributions of individual monomers obtained from HyRes (**g**) and all-atom simulations (**h**). Monomers 10 and 30 were randomly selected to highlight the difference in convergence of the HyRes and all-atom simulations.

**Extended Data Fig. 9.**
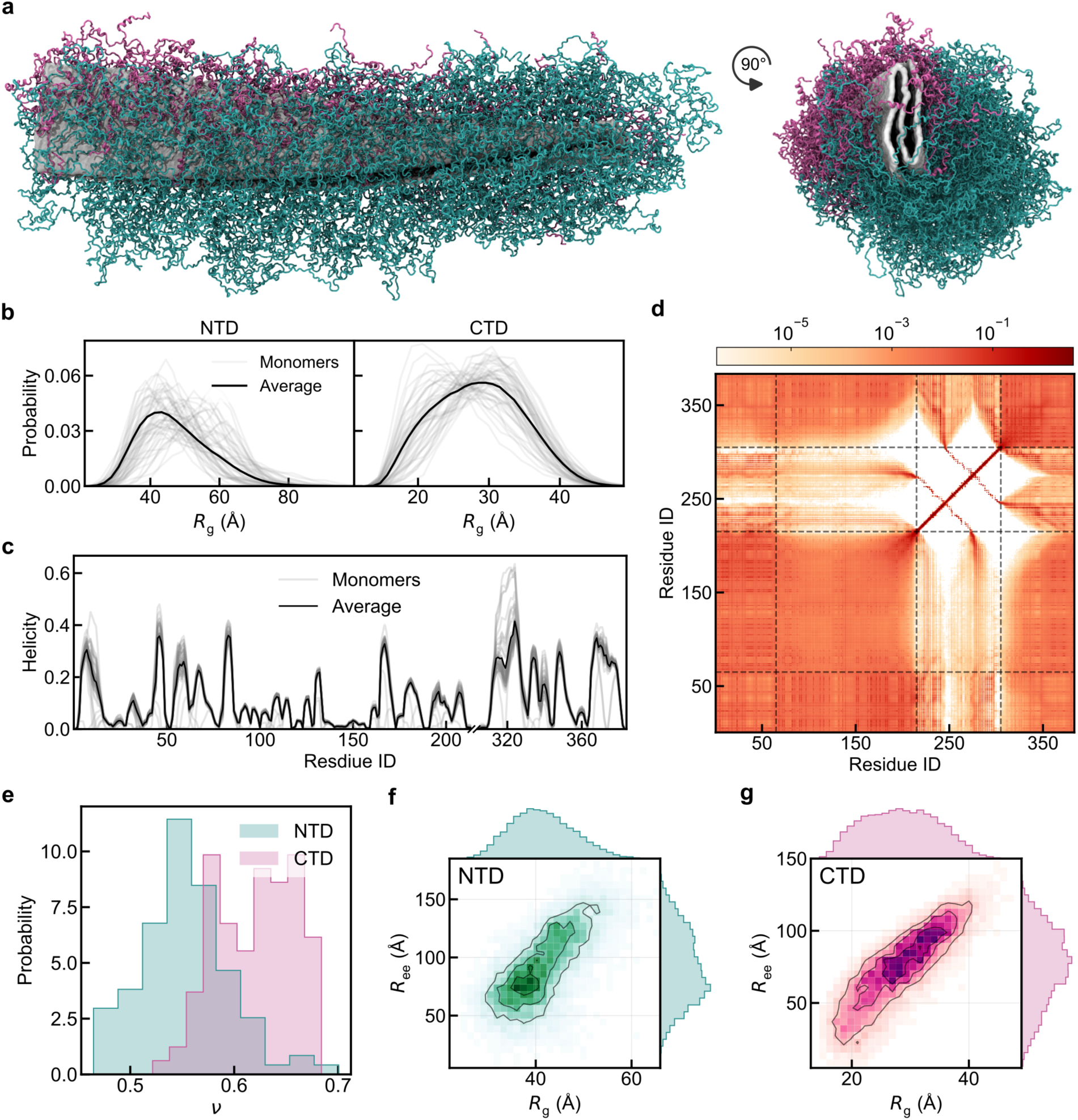
HyRes simulation of full-length tau fibril. **a**, A representative snapshot of the full-length tau fibril containing the structured core (grey surfaces), disordered N-terminal (teal coil chains), and C-terminal regions (magenta coils). **b-c**, Individual and average distributions of *R*_g_ (**b**, for the NTD and CTD, separately) and residual helicity (**c**) for the central 50 monomers. **d**, Probability (in log scale) of residue-residue contacts among tau monomers within the full-length fibril. **e**, Distributions of scaling exponents, *v*, for the NTD and CTD. f-g, Joint distribution of *R*_g_ and *R*_ee_ for the NTD (**f**) and CTD (**g**), with marginal distributions displayed on the top and right.

**Extended Data Fig. 10.**
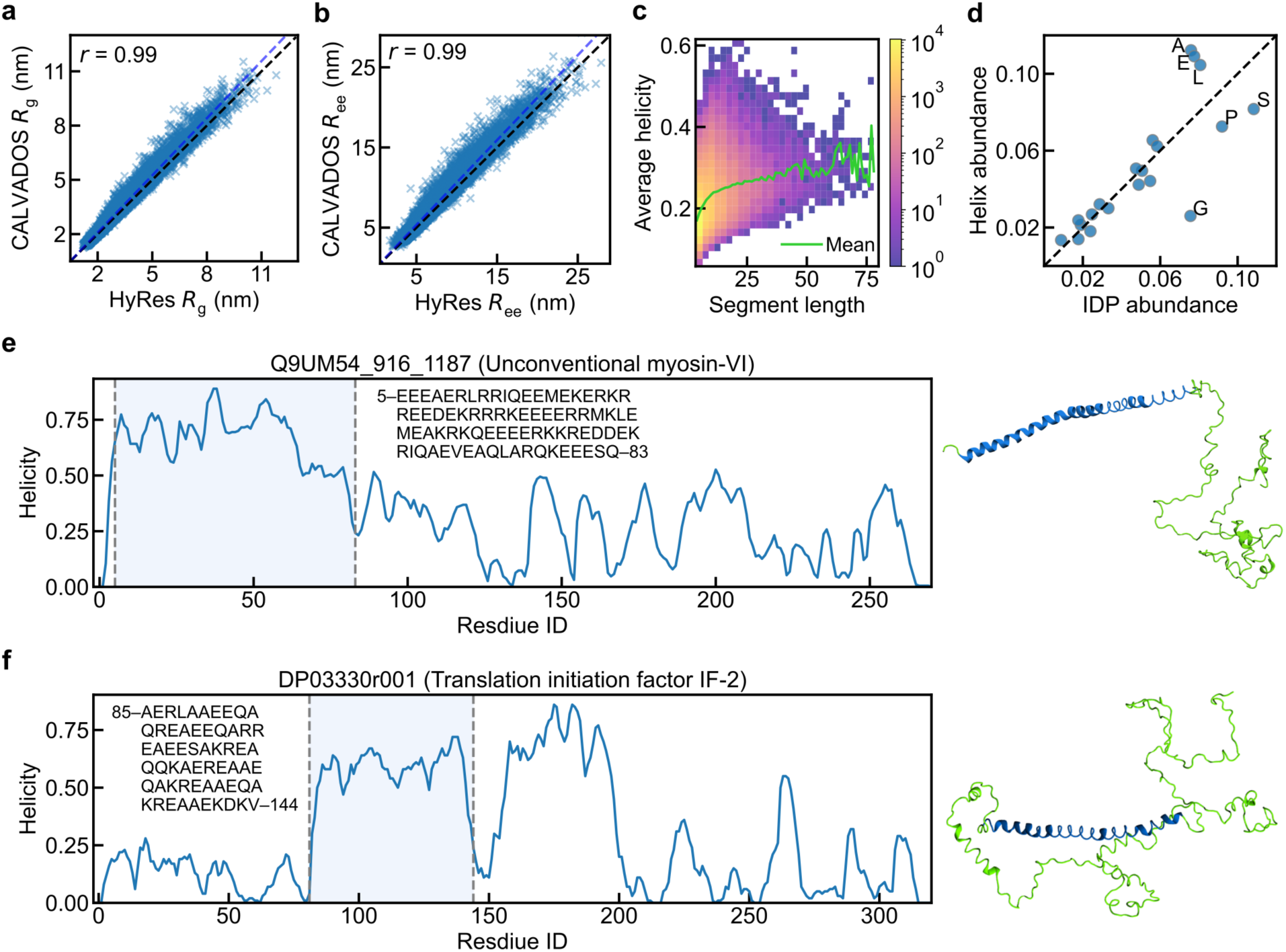
General features of HyRes-IDRome. **a-b**, Correlations between HyRes-and CALVADOS-derived *R*_g_ (**a**) and *R*_ee_ (**b**) for IDRome. **c**, For transient helical segments, the relation between segment length and the helicity. The green curve shows the correlation between the average residual helicity of same-length segments and segment length. **d**, The abundance of each amino acid type in the whole IDRome (IDP abundance) and all helical segments (helix abundance). **e-f**, Helicity profiles and snapshots of representative long helical segments, whose sequences (as well as the starting and ending residue indices) are shown as inserts.

## Reference

1. Tyson, J. J., Chen, K. & Novak, B. Network dynamics and cell physiology. Nat Rev Mol Cell Biol 2, 908–916 (2001).

2. Wei, G., Xi, W., Nussinov, R. & Ma, B. Protein Ensembles: How Does Nature Harness Thermodynamic Fluctuations for Life? The Diverse Functional Roles of Conformational Ensembles in the Cell. Chem. Rev. 116, 6516–6551 (2016).

3. Wright, P. E. & Dyson, H. J. Intrinsically disordered proteins in cellular signalling and regulation. Nat Rev Mol Cell Biol 16, 18–29 (2015).

4. Hatos, A. et al. DisProt: intrinsic protein disorder annotation in 2020. Nucleic Acids Res 48, D269–D276 (2020).

5. Ruff, K. M. et al. Molecular grammars of predicted intrinsically disordered regions that span the human proteome. Cell 189, 323–342.e17 (2026).

6. Iakoucheva, L. M., Brown, C. J., Lawson, J. D., Obradović, Z. & Dunker, A. K. Intrinsic Disorder in Cell-signaling and Cancer-associated Proteins. Journal of Molecular Biology 323, 573–584 (2002).

7. Ruff, K. M., Pappu, R. V. & Holehouse, A. S. Conformational preferences and phase behavior of intrinsically disordered low complexity sequences: insights from multiscale simulations. Current Opinion in Structural Biology 56, 1–10 (2019).

8. Liu, Z. H., Tsanai, M., Zhang, O., Head-Gordon, T. & Forman-Kay, J. D. Biological insights from integrative modeling of intrinsically disordered protein systems. Current Opinion in Structural Biology 93, 103063 (2025).

9. Wu, H. & Fuxreiter, M. The Structure and Dynamics of Higher-Order Assemblies: Amyloids, Signalosomes, and Granules. Cell 165, 1055–1066 (2016).

10. Borgia, A. et al. Extreme disorder in an ultrahigh-affinity protein complex. Nature 555, 61–66 (2018).

11. Bugge, K. et al. Role of charges in a dynamic disordered complex between an IDP and a folded domain. Nat Commun 16, 3242 (2025).

12. Chen, J., Liu, X. & Chen, J. Targeting Intrinsically Disordered Proteins through Dynamic Interactions. Biomolecules 10, 743 (2020).

13. Brangwynne, C. P. et al. Germline P Granules Are Liquid Droplets That Localize by Controlled Dissolution/Condensation. Science 324, 1729–1732 (2009).

14. Li, P. et al. Phase transitions in the assembly of multivalent signalling proteins. Nature 483, 336–340 (2012).

15. Brangwynne, C. P., Tompa, P. & Pappu, R. V. Polymer physics of intracellular phase transitions. Nature Phys 11, 899–904 (2015).

16. Kilgore, H. R., Moreno, S. & Young, R. A. Protein codes and mobility together shape cellular function and disease. Trends in Biochemical Sciences 51, 8–26 (2026).

17. Uversky, V. N., Oldfield, C. J. & Dunker, A. K. Intrinsically Disordered Proteins in Human Diseases: Introducing the D2 Concept. Annual Review of Biophysics 37, 215–246 (2008).

18. Tsafou, K., Tiwari, P. B., Forman-Kay, J. D., Metallo, S. J. & Toretsky, J. A. Targeting Intrinsically Disordered Transcription Factors: Changing the Paradigm. Journal of Molecular Biology 430, 2321–2341 (2018).

19. Shirnekhi, H. K., Chandra, B. & Kriwacki, R. W. The Role of Phase-Separated Condensates in Fusion Oncoprotein–Driven Cancers. Annual Review of Cancer Biology 7, 73–91 (2023).

20. Ghafouri, H. et al. Toward a unified framework for determining conformational ensembles of disordered proteins. Nat Methods 23, 705–719 (2026).

21. Berlow, R. B., Dyson, H. J. & Wright, P. E. Expanding the Paradigm: Intrinsically Disordered Proteins and Allosteric Regulation. Journal of Molecular Biology 430, 2309–2320 (2018).

22. Camacho-Zarco, A. R. et al. NMR Provides Unique Insight into the Functional Dynamics and Interactions of Intrinsically Disordered Proteins. Chem. Rev. 122, 9331–9356 (2022).

23. Koch, M. H. J., Vachette, P. & Svergun, D. I. Small-angle scattering: a view on the properties, structures and structural changes of biological macromolecules in solution. Quarterly Reviews of Biophysics 36, 147–227 (2003).

24. Schuler, B., Soranno, A., Hofmann, H. & Nettels, D. Single-Molecule FRET Spectroscopy and the Polymer Physics of Unfolded and Intrinsically Disordered Proteins. Annual Review of Biophysics 45, 207–231 (2016).

25. Stuchfield, D. & Barran, P. Unique insights to intrinsically disordered proteins provided by ion mobility mass spectrometry. Current Opinion in Chemical Biology 42, 177–185 (2018).

26. Balasubramaniam, D. & Komives, E. A. Hydrogen-exchange mass spectrometry for the study of intrinsic disorder in proteins. Biochimica et Biophysica Acta (BBA) - Proteins and Proteomics 1834, 1202–1209 (2013).

27. Chen, J. Towards the physical basis of how intrinsic disorder mediates protein function. Archives of Biochemistry and Biophysics 524, 123–131 (2012).

28. Bonomi, M., Heller, G. T., Camilloni, C. & Vendruscolo, M. Principles of protein structural ensemble determination. Current Opinion in Structural Biology 42, 106–116 (2017).

29. Shea, J.-E., Best, R. B. & Mittal, J. Physics-based computational and theoretical approaches to intrinsically disordered proteins. Current Opinion in Structural Biology 67, 219–225 (2021).

30. Tesei, G. et al. Conformational ensembles of the human intrinsically disordered proteome. Nature 626, 897–904 (2024).

31. Wang, L., Brasnett, C., Borges-Araújo, L., Souza, P. C. T. & Marrink, S. J. Martini3-IDP: improved Martini 3 force field for disordered proteins. Nat Commun 16, 2874 (2025).

32. Vitalis, A. & Pappu, R. V. ABSINTH: A new continuum solvation model for simulations of polypeptides in aqueous solutions. Journal of Computational Chemistry 30, 673–699 (2009).

33. Best, R. B., Zheng, W. & Mittal, J. Balanced Protein–Water Interactions Improve Properties of Disordered Proteins and Non-Specific Protein Association. J. Chem. Theory Comput. 10, 5113–5124 (2014).

34. Huang, J. et al. CHARMM36m: an improved force field for folded and intrinsically disordered proteins. Nat Methods 14, 71–73 (2017).

35. Robustelli, P., Piana, S. & Shaw, D. E. Developing a molecular dynamics force field for both folded and disordered protein states. Proceedings of the National Academy of Sciences 115, E4758–E4766 (2018).

36. Gong, X., Zhang, Y. & Chen, J. Advanced Sampling Methods for Multiscale Simulation of Disordered Proteins and Dynamic Interactions. Biomolecules 11, 1416 (2021).

37. Bottaro, S. & Lindorff-Larsen, K. Biophysical experiments and biomolecular simulations: A perfect match? Science 361, 355–360 (2018).

38. Chen, J., Im, W. & Brooks, C. L. Balancing Solvation and Intramolecular Interactions: Toward a Consistent Generalized Born Force Field. J. Am. Chem. Soc. 128, 3728–3736 (2006).

39. Choi, J.-M. & Pappu, R. V. Improvements to the ABSINTH Force Field for Proteins Based on Experimentally Derived Amino Acid Specific Backbone Conformational Statistics. J. Chem. Theory Comput. 15, 1367–1382 (2019).

40. Maffucci, I. & Contini, A. An Updated Test of AMBER Force Fields and Implicit Solvent Models in Predicting the Secondary Structure of Helical, β-Hairpin, and Intrinsically Disordered Peptides. J. Chem. Theory Comput. 12, 714–727 (2016).

41. Regy, R. M., Thompson, J., Kim, Y. C. & Mittal, J. Improved coarse-grained model for studying sequence dependent phase separation of disordered proteins. Protein Science 30, 1371–1379 (2021).

42. Latham, A. P. & Zhang, B. Consistent Force Field Captures Homologue-Resolved HP1 Phase Separation. J. Chem. Theory Comput. 17, 3134–3144 (2021).

43. Tesei, G., Schulze, T. K., Crehuet, R. & Lindorff-Larsen, K. Accurate model of liquid–liquid phase behavior of intrinsically disordered proteins from optimization of single-chain properties. Proceedings of the National Academy of Sciences 118, e2111696118 (2021).

44. Wu, H., Wolynes, P. G. & Papoian, G. A. AWSEM-IDP: A Coarse-Grained Force Field for Intrinsically Disordered Proteins. J. Phys. Chem. B 122, 11115–11125 (2018).

45. Valdes-Garcia, G., Heo, L., Lapidus, L. J. & Feig, M. Modeling Concentration-dependent Phase Separation Processes Involving Peptides and RNA via Residue-Based Coarse-Graining. J. Chem. Theory Comput. 19, 669–678 (2023).

46. Gianni, S. et al. Fuzziness and Frustration in the Energy Landscape of Protein Folding, Function, and Assembly. Acc. Chem. Res. 54, 1251–1259 (2021).

47. Janson, G. & Feig, M. Transferable deep generative modeling of intrinsically disordered protein conformations. PLOS Computational Biology 20, e1012144 (2024).

48. Zhu, J. et al. Accurate Generation of Conformational Ensembles for Intrinsically Disordered Proteins with IDPFold. Advanced Science 12, e11636 (2025).

49. Novak, B., Lotthammer, J. M., Emenecker, R. J. & Holehouse, A. S. Accurate predictions of disordered protein ensembles with STARLING. Nature 1–11 (2026) doi:10.1038/s41586-026-10141-2.

50. Lewis, S. et al. Scalable emulation of protein equilibrium ensembles with generative deep learning. Science 389, eadv9817 (2025).

51. Liu, X. & Chen, J. HyRes: a coarse-grained model for multi-scale enhanced sampling of disordered protein conformations. Physical Chemistry Chemical Physics 19, 32421–32432 (2017).

52. Zhang, Y., Liu, X. & Chen, J. Toward Accurate Coarse-Grained Simulations of Disordered Proteins and Their Dynamic Interactions. J. Chem. Inf. Model. 62, 4523–4536 (2022).

53. Zhang, Y., Li, S., Gong, X. & Chen, J. Toward Accurate Simulation of Coupling between Protein Secondary Structure and Phase Separation. J. Am. Chem. Soc. 146, 342–357 (2024).

54. Shorkey, S. A. et al. Tracking flaviviral protease conformational dynamics by tuning single-molecule nanopore tweezers. Biophysical Journal 124, 145–157 (2025).

55. Zhao, J. et al. Intrinsically Disordered N-terminal Domain (NTD) of p53 Interacts with Mitochondrial PTP Regulator Cyclophilin D. Journal of Molecular Biology 434, 167552 (2022).

56. Barethiya, S., Schultz, S., Zhang, Y. & Chen, J. Coarse-Grained Simulations of Phosphorylation Regulation of p53 Autoinhibition. Biochemistry 64, 1636–1645 (2025).

57. Das, R. K. & Pappu, R. V. Conformations of intrinsically disordered proteins are influenced by linear sequence distributions of oppositely charged residues. Proceedings of the National Academy of Sciences 110, 13392–13397 (2013).

58. Shi, W. H., Adhikari, R. S., Asthagiri, D. N. & Marciel, A. B. Influence of Charge Block Length on Conformation and Solution Behavior of Polyampholytes. ACS Macro Lett. 12, 195–200 (2023).

59. Tesei, G. et al. Self-association of a highly charged arginine-rich cell-penetrating peptide. Proceedings of the National Academy of Sciences 114, 11428–11433 (2017).

60. Bremer, A. et al. Deciphering how naturally occurring sequence features impact the phase behaviours of disordered prion-like domains. Nat. Chem. 14, 196–207 (2022).

61. Tesei, G. & Lindorff-Larsen, K. Improved predictions of phase behaviour of intrinsically disordered proteins by tuning the interaction range. Open Res Europe 2, 94 (2023).

62. Holla, A. et al. Identifying Sequence Effects on Chain Dimensions of Disordered Proteins by Integrating Experiments and Simulations. JACS Au 4, 4729–4743 (2024).

63. Liu, X. & Chen, J. Residual Structures and Transient Long-Range Interactions of p53 Transactivation Domain: Assessment of Explicit Solvent Protein Force Fields. J. Chem. Theory Comput. 15, 4708–4720 (2019).

64. Bertoncini, C. W. et al. Release of long-range tertiary interactions potentiates aggregation of natively unstructured α-synuclein. Proceedings of the National Academy of Sciences 102, 1430–1435 (2005).

65. Tesei, G. et al. DEER-PREdict: Software for efficient calculation of spin-labeling EPR and NMR data from conformational ensembles. PLOS Computational Biology 17, e1008551 (2021).

66. Marsh, J. A., Singh, V. K., Jia, Z. & Forman-Kay, J. D. Sensitivity of secondary structure propensities to sequence differences between α- and γ-synuclein: Implications for fibrillation. Protein Science 15, 2795–2804 (2006).

67. Heo, L. & Feig, M. One bead per residue can describe all-atom protein structures. Structure 32, 97–111.e6 (2024).

68. Han, B., Liu, Y., Ginzinger, S. W. & Wishart, D. S. SHIFTX2: significantly improved protein chemical shift prediction. J Biomol NMR 50, 43–57 (2011).

69. Hoch, J. C. et al. Biological Magnetic Resonance Data Bank. Nucleic Acids Res 51, D368–D376 (2023).

70. Kabsch, W. & Sander, C. Dictionary of protein secondary structure: Pattern recognition of hydrogen-bonded and geometrical features. Biopolymers 22, 2577–2637 (1983).

71. Kjaergaard, M. & Poulsen, F. M. Sequence correction of random coil chemical shifts: correlation between neighbor correction factors and changes in the Ramachandran distribution. J Biomol NMR 50, 157–165 (2011).

72. Galvanetto, N. et al. Extreme dynamics in a biomolecular condensate. Nature 619, 876–883 (2023).

73. Zhang, J. Y., Dregni, A. J. & Hong, M. Heterogeneous Dynamics of the Fuzzy Coat of Full-Length Phospho-Mimetic Tau Fibrils. J. Am. Chem. Soc. 148, 1623–1637 (2026).

74. Wang, Y. & Mandelkow, E. Tau in physiology and pathology. Nat Rev Neurosci 17, 22–35 (2016).

75. Pacheco, S. et al. Phosphorylation of α-Synuclein Fibrils at S129 Changes DNAJB1 Binding as Probed by Solid-State NMR. JACS Au 6, 343–356 (2026).

76. Faidon Brotzakis, Z., et al. Determination of the Structure and Dynamics of the Fuzzy Coat of an Amyloid Fibril of IAPP Using Cryo-Electron Microscopy. Biochemistry 62, 2407–2416 (2023).

77. Nguyen, P. H. et al. Amyloid Oligomers: A Joint Experimental/Computational Perspective on Alzheimer’s Disease, Parkinson’s Disease, Type II Diabetes, and Amyotrophic Lateral Sclerosis. Chem. Rev. 121, 2545–2647 (2021).

78. Joseph, J. A. et al. Physics-driven coarse-grained model for biomolecular phase separation with near-quantitative accuracy. Nat Comput Sci 1, 732–743 (2021).

79. Valentini, E., Kikhney, A. G., Previtali, G., Jeffries, C. M. & Svergun, D. I. SASBDB, a repository for biological small-angle scattering data. Nucleic Acids Res 43, D357–D363 (2015).

80. Eastman, P. et al. OpenMM 8: Molecular Dynamics Simulation with Machine Learning Potentials. J. Phys. Chem. B 128, 109–116 (2024).

81. Gowers, R. J. et al. MDAnalysis: A Python Package for the Rapid Analysis of Molecular Dynamics Simulations. SciPy 2016 https://doi.org/10.25080/Majora-629e541a-00e (2016) doi:10.25080/Majora-629e541a-00e.

82. Michaud-Agrawal, N., Denning, E. J., Woolf, T. B. & Beckstein, O. MDAnalysis: A toolkit for the analysis of molecular dynamics simulations. Journal of Computational Chemistry 32, 2319–2327 (2011).

83. Abraham, M. J. et al. GROMACS: High performance molecular simulations through multi-level parallelism from laptops to supercomputers. SoftwareX 1–2, 19–25 (2015).

84. Gartner, T. E. & Jayaraman, A. Modeling and Simulations of Polymers: A Roadmap. Macromolecules 52, 755–786 (2019).

85. Martinsen, J. H. et al. Structure, dynamics, and stability of the globular domain of human linker histone H1.0 and the role of positive charges. Protein Science 31, 918–932 (2022).

86. Kumar, S., Rosenberg, J. M., Bouzida, D., Swendsen, R. H. & Kollman, P. A. THE weighted histogram analysis method for free-energy calculations on biomolecules. I. The method. Journal of Computational Chemistry 13, 1011–1021 (1992).

87. Lab, G. WHAM: an implementation of the weighted histogram analysis method. (2026).

88. Webb, B. & Sali, A. Comparative Protein Structure Modeling Using MODELLER. Current Protocols in Bioinformatics 54, 5.6.1–5.6.37 (2016).

89. Mészáros, B., Erdős, G. & Dosztányi, Z. IUPred2A: context-dependent prediction of protein disorder as a function of redox state and protein binding. Nucleic Acids Res 46, W329–W337 (2018).

90. Holehouse, A. S., Ahad, J., Das, R. K. & Pappu, R. V. CIDER: Classification of Intrinsically Disordered Ensemble Regions. Biophysical Journal 108, 228a (2015).

91. Huerta-Cepas, J., Serra, F. & Bork, P. ETE 3: Reconstruction, Analysis, and Visualization of Phylogenomic Data. Mol Biol Evol 33, 1635–1638 (2016).

