## Supplemental tables and figures for "HyRes: Accurate Physics-Based Simulation of Dynamic Protein Structures and Interactions in Complex Environments at Scale"

### Supplementary Tables

**Supplementary Table S1.** $R_g$ of 20 A1-LCD variants from experiments[1] and simulations

| A1-LCD variants | Experimental $R_g$ (Å) | HyRes $R_g$ (Å) |
|---|---|---|
| A1-NLS | 27.6 | 26.5 |
| A1+NLS | 25.8 | 26.0 |
| A1-10R | 26.7 | 26.4 |
| A1-6R | 25.7 | 25.9 |
| A1+2R | 26.2 | 26.4 |
| A1+7R | 27.1 | 27.1 |
| A1-3R+3K | 26.3 | 27.3 |
| A1-6R+6K | 27.9 | 28.0 |
| A1-10R+10K | 28.5 | 28.9 |
| A1+12D | 28.0 | 28.2 |
| A1+4D | 27.2 | 26.4 |
| A1+8D | 26.9 | 27.7 |
| A1-9F+3Y | 26.8 | 27.6 |
| A1+12E | 28.5 | 28.6 |
| A1+7K+12D | 29.2 | 30.0 |
| A1-4D | 26.4 | 26.2 |
| A1-8F+4Y | 27.1 | 26.9 |
| A1+7F-7Y | 27.2 | 27.3 |
| A1-12F+12Y | 26.0 | 25.7 |
| A1-9F+6Y | 26.5 | 26.8 |

**Supplementary Table S2.** $R_g$ of 30 EK25 IDPs from ABSINTH[2] and HyRes simulations

| EK25 variants | κ | ABSINTH $R_g$ (Å) | HyRes $R_g$ (Å) ($\varepsilon_r$ = 20.0) | HyRes $R_g$ (Å) ($\varepsilon_r$ = 60.0) |
|---|---|---|---|---|
| sv1 | 0.0009 | 29.9 | 13.8 | 21.1 |
| sv2 | 0.0025 | 30.2 | 14.6 | 24.0 |
| sv3 | 0.0139 | 27.1 | 13.9 | 20.2 |
| sv4 | 0.0140 | 28.7 | 14.2 | 21.6 |
| sv5 | 0.0245 | 28.5 | 14.4 | 21.1 |
| sv6 | 0.0273 | 28.3 | 14.3 | 22.3 |
| sv7 | 0.0450 | 28.7 | 14.5 | 22.1 |
| sv8 | 0.0450 | 28.8 | 14.5 | 21.7 |
| sv9 | 0.0624 | 24.9 | 13.4 | 17.5 |
| sv10 | 0.0834 | 25.5 | 13.8 | 17.8 |
| sv11 | 0.0841 | 27.0 | 13.8 | 20.1 |
| sv12 | 0.0864 | 22.1 | 13.3 | 16.5 |
| sv13 | 0.0951 | 24.1 | 13.4 | 16.1 |
| sv14 | 0.1311 | 25.3 | 13.5 | 18.2 |
| sv15 | 0.1354 | 20.4 | 13.1 | 15.4 |
| sv16 | 0.1458 | 25.1 | 13.6 | 17.0 |
| sv17 | 0.1643 | 20.2 | 13.1 | 15.6 |
| sv18 | 0.1677 | 26.6 | 13.4 | 18.7 |
| sv19 | 0.1941 | 27.5 | 14.2 | 20.1 |
| sv20 | 0.2721 | 19.6 | 13.2 | 15.7 |
| sv21 | 0.2737 | 22.5 | 12.8 | 15.5 |
| sv22 | 0.3218 | 20.5 | 12.8 | 15.1 |
| sv23 | 0.3545 | 19.3 | 12.9 | 15.2 |
| sv24 | 0.4456 | 17.6 | 12.3 | 14.1 |
| sv25 | 0.5283 | 19.5 | 13.3 | 15.5 |
| sv26 | 0.6101 | 17.5 | 12.4 | 13.9 |
| sv27 | 0.6729 | 19.6 | 12.6 | 14.7 |
| sv28 | 0.7666 | 17.9 | 12.8 | 14.0 |
| sv29 | 0.8764 | 17.0 | 12.3 | 13.6 |
| sv30 | 1.0000 | 16.8 | 11.9 | 13.3 |

**Supplementary Table S3.** $R_g$ of 98 IDPs from experiments and simulations. The first 41 IDPs are collected from ref 2[3] and the others are from SASBDB showing their IDs

| IDPs | Exp. $R_g$ (Å) | Sim. $R_g$ (Å) | IDPs | Exp. $R_g$ (Å) | Sim. $R_g$ (Å) |
|---|---|---|---|---|---|
| Hst5 | 13.8 | 12.8 | SASDEV2 | 25.0 | 24.0 |
| Hst52 | 18.7 | 20.2 | SASDEZ2 | 30.0 | 30.3 |
| p53 | 28.7 | 29.1 | SASDC53 | 28.0 | 26.4 |
| ACTR | 26.0 | 24.6 | SASDEB3 | 29.0 | 32.1 |
| Ash1 | 29.0 | 27.5 | SASDEC3 | 41.0 | 38.4 |
| CTD2 | 26.1 | 23.5 | SASDSC3 | 11.0 | 10.3 |
| Sic1 | 30.0 | 28.0 | SASDSE3 | 10.0 | 11.6 |
| SH4UD | 29.0 | 23.8 | SASDSG3 | 12.0 | 11.4 |
| p15PAF | 28.0 | 31.5 | SASDSH3 | 13.0 | 11.5 |
| ProTa | 37.9 | 40.6 | SASDSJ3 | 11.0 | 10.0 |
| hNL3cyt | 32.0 | 30.5 | SASDS74 | 13.0 | 12.3 |
| RNaseA | 34.0 | 33.1 | SASDSH4 | 12.0 | 12.7 |
| hNHE1 | 36.3 | 34.1 | SASDF34 | 47.0 | 47.0 |
| aSyn | 35.5 | 34.8 | SASDLS4 | 46.0 | 52.4 |
| FhuA | 33.4 | 33.4 | SASDLT4 | 63.0 | 62.6 |
| TDP43 | 28.0 | 29.2 | SASDLU4 | 67.0 | 65.2 |
| K18 | 38.0 | 34.6 | SASDEX4 | 24.0 | 26.2 |
| FUS | 33.2 | 31.7 | SASDJB5 | 62.0 | 62.9 |
| K27 | 37.0 | 42.2 | SASDLM5 | 21.0 | 27.1 |
| K10 | 40.0 | 39.5 | SASDBY5 | 51.0 | 43.3 |
| LAF-1 | 30.8 | 31.2 | SASDJ86 | 35.0 | 41.2 |
| K25 | 41.0 | 38.6 | SASDDF6 | 33.0 | 37.1 |
| K32 | 42.0 | 45.3 | SASDJJ6 | 35.0 | 36.9 |
| CAHSD | 48.0 | 46.5 | SASDJK6 | 39.0 | 36.4 |
| Ddx4 | 36.1 | 38.2 | SASDVN9 | 35.0 | 36.7 |
| K23 | 49.0 | 48.2 | SASDU37 | 36.0 | 35.7 |
| K44 | 52.0 | 50.5 | SASDCY9 | 23.0 | 30.6 |
| PNt | 51.0 | 46.9 | SASDHH8 | 15.0 | 13.1 |
| PNtSwap1 | 49.0 | 47.5 | SASDL99 | 42.0 | 41.8 |
| PNtSwap4 | 53.0 | 47.2 | SASDL89 | 65.0 | 54.7 |
| PNtSwap5 | 49.0 | 43.3 | SASDL79 | 43.0 | 39.2 |
| PNtSwap6 | 53.0 | 47.7 | SASDL69 | 53.0 | 52.1 |

| | | | | | |
|---|---|---|---|---|---|
| GHRICD | 60.0 | 60.8 | SASDJS7 | 35.0 | 39.3 |
| DSS1 | 25.0 | 27.1 | SASDK68 | 37.0 | 38.0 |
| p27Cv14 | 29.4 | 31.0 | SASDKT8 | 31.0 | 27.7 |
| p27Cv15 | 29.2 | 31.2 | SASDFK8 | 31.0 | 31.7 |
| p27Cv31 | 28.1 | 30.3 | SASDEF6 | 27.0 | 24.5 |
| p27Cv44 | 24.9 | 27.0 | SASDP88 | 52.0 | 47.2 |
| p27Cv56 | 23.3 | 23.6 | SASDQM8 | 32.0 | 29.1 |
| p27Cv78 | 22.1 | 22.1 | SASDQN8 | 33.0 | 29.8 |
| NHE6cmdd | 32.0 | 29.7 | SASDLF9 | 34.0 | 33.2 |
| SASDC62 | 17.0 | 17.1 | SASDQC7 | 40.0 | 33.5 |
| SASDD92 | 67.0 | 68.8 | SASDQD7 | 28.0 | 21.8 |
| SASDED2 | 37.0 | 43.7 | SASDQE7 | 27.0 | 22.3 |
| SASDEE2 | 30.0 | 31.5 | SASDQF7 | 37.0 | 34.1 |
| SASDEH2 | 16.0 | 17.0 | SASDTZ6 | 47.0 | 39.5 |
| SASDEM2 | 24.0 | 19.8 | SASDU67 | 34.0 | 35.1 |
| SASDSN2 | 23.0 | 24.6 | SASDKD6 | 25.0 | 24.9 |
| SASDER2 | 32.0 | 30.6 | SASDKH8 | 38.0 | 36.7 |

**Supplementary Table S4.** smFRET-derived dye pair distances and simulated Cα pair distances of 16 IDPs[4]

| IDPs | Alexa 488/594 distance (Å) | Cy3B/CF660R distance (Å) | Simulated Cα distance (Å) |
|---|---|---|---|
| sGrich | 40.0 | 32.0 | 44.5 |
| dkh | 47.0 | 44.0 | 45.9 |
| dGrich | 51.0 | 46.0 | 52.2 |
| sPTBP | 55.0 | 49.0 | 49.0 |
| sCh | 57.0 | 53.0 | 55.7 |
| sNrich | 59.0 | 53.0 | 57.5 |
| skh | 59.0 | 47.0 | 55.1 |
| dCh+ | 59.0 | 47.0 | 52.3 |
| dCh- | 67.0 | 62.0 | 58.5 |
| skl | 67.0 | 59.0 | 62.0 |
| sNh+ | 67.0 | 53.0 | 57.6 |
| dArich | 68.0 | 64.0 | 55.2 |
| dErich | 69.0 | 64.0 | 58.0 |
| dTRBP | 71.0 | 65.0 | 55.8 |
| dkl | 72.0 | 67.0 | 59.2 |
| sNh- | 73.0 | 71.0 | 64.6 |

**Supplementary Table S5.** Pearson correlation (*r*) and RMSD of HyRes-derived PRE profiles. The spin-labeling sites measured for each IDP are also listed.

| IDPs | Label sites | Pearson *r* | RMSD |
|---|---|---|---|
| aSyn[5–7] | 18, 24, 42, 62, 76, 87, 90, 103, 140 | 0.76 ± 0.19 | 0.19 ±0.04 |
| ACTR[8] | 3, 21, 41, 61 | 0.60 ± 0.09 | 0.24 ± 0.03 |
| Osteopontin[9] | 10, 64, 144, 203 | 0.69 ± 0.05 | 0.19 ± 0.02 |
| Darpp-32[10] | 7, 38, 53, 74, 96 | 0.64 ± 0.11 | 0.22 ± 0.01 |
| RNAPd[11] | 110, 132, 151, 168 | 0.80 ± 0.06 | 0.16 ± 0.04 |
| ER-NTD[12] | 10, 46, 84, 106, 137, 167 | 0.59 ± 0.12 | 0.21 ± 0.02 |
| Tau[13] | 15, 72, 125, 178, 239, 256, 322, 352, 384, 416 | 0.79 ± 0.07 | 0.14 ± 0.02 |
| K18[14] | 17, 48, 79, 111 | 0.86 ± 0.03 | 0.19 ± 0.03 |
| p53TAD[15] | 7, 28, 39, 61 | 0.75 ± 0.08 | 0.17 ± 0.06 |

**Supplementary Table S6.** RMSD of PRE profiles predicted by various all-atom force fields and HyRes

| **Force fields** | **RMSD** | | |
|---|---|---|---|
| | aSyn[16] | ACTR[16] | p53TAD[17] |
| HyRes | 0.19 | 0.24 | 0.17 |
| a99SB-disp | 0.17 | 0.22 | 0.27 |
| a99SB-ILDN TIP4P-D | 0.18 | 0.23 | 0.27 |
| a99SBUCB | 0.17 | 0.20 | - |
| a03ws | 0.17 | 0.24 | - |
| C36m | 0.32 | 0.18 | 0.36 |
| C22* TIP3P | 0.35 | 0.20 | 0.49 |
| a99SB-ILDN TIP3P | 0.39 | 0.38 | 0.56 |

**Supplementary Table S7.** Pearson correlation and RMSD (ppm) of HyRes-predicted chemical shifts and secondary shifts (Δδ) of 58 IDPs

| BMRB ID | Pearson correlation/RMSD | | | | | | |
|---|---|---|---|---|---|---|---|
| | Cα | Hα | C’ | Cβ | N | HN | Δδ |
| 6857 | 0.99/0.65 | 0.94/0.13 | — | 1.00/1.02 | — | 0.73/0.25 | 0.32/1.02 |
| 15123 | 0.99/0.70 | 0.86/0.11 | — | 1.00/0.27 | 0.96/1.12 | 0.58/0.17 | 0.69/0.85 |
| 15141 | 0.98/0.73 | — | 0.86/0.76 | 1.00/0.38 | 0.90/1.58 | 0.50/0.18 | 0.43/0.97 |
| 15397 | 1.00/0.39 | — | 0.95/0.42 | 1.00/0.24 | 0.94/1.51 | 0.64/0.14 | 0.34/0.59 |
| 15767 | 0.99/0.68 | — | 0.86/0.62 | 1.00/0.45 | 0.96/2.75 | 0.34/0.33 | 0.28/0.91 |
| 16165 | 0.99/0.66 | — | 0.92/0.47 | 1.00/0.32 | 0.92/1.55 | 0.25/0.27 | 0.39/0.81 |
| 16300 | 1.00/0.95 | — | 0.91/0.58 | 1.00/0.94 | 0.96/1.74 | 0.53/0.24 | 0.46/0.57 |
| 16657 | 1.00/1.02 | 0.90/0.17 | — | 1.00/0.52 | 0.93/1.52 | 0.29/0.18 | 0.62/0.76 |
| 16798 | 1.00/0.45 | — | 0.95/0.58 | 1.00/0.29 | 0.98/1.20 | 0.53/0.18 | 0.51/0.60 |
| 18198 | 1.00/0.46 | — | — | 1.00/0.24 | 0.94/1.51 | 0.49/0.17 | 0.43/0.58 |
| 18841 | 0.97/0.74 | 0.86/0.12 | 0.89/0.72 | 1.00/0.37 | 0.86/1.34 | 0.60/0.15 | 0.63/1.00 |
| 19332 | 1.00/0.48 | 0.95/0.06 | 0.91/0.57 | 1.00/0.31 | 0.94/1.60 | 0.31/0.18 | 0.48/0.67 |
| 19830 | 1.00/3.24 | 0.95/0.11 | 0.93/2.69 | 1.00/3.17 | 0.94/1.54 | 0.50/0.16 | 0.55/0.68 |
| 25384 | 0.99/0.55 | 0.91/0.08 | 0.93/0.57 | 1.00/0.54 | 0.92/1.61 | 0.41/0.23 | 0.18/0.82 |
| 26549 | 0.99/0.56 | 0.91/0.07 | 0.95/0.48 | 1.00/0.33 | 0.95/1.47 | 0.40/0.19 | 0.57/0.70 |
| 26639 | 1.00/0.61 | — | 0.93/0.54 | 1.00/0.31 | 0.96/1.11 | 0.45/0.16 | 0.66/0.64 |
| 26662 | 0.99/1.03 | — | 0.78/0.90 | 1.00/0.36 | 0.86/1.62 | 0.39/0.21 | 0.48/1.31 |
| 26672 | 1.00/0.45 | — | 0.85/0.65 | 1.00/0.27 | 0.97/1.14 | 0.44/0.13 | 0.41/0.44 |
| 26719 | 0.99/0.53 | — | 0.90/0.74 | 1.00/0.36 | 0.92/1.48 | 0.25/0.17 | 0.43/0.56 |
| 26762 | 1.00/0.67 | — | 0.95/0.43 | 1.00/0.36 | 0.95/1.76 | 0.46/0.35 | 0.38/0.88 |
| 26823 | 1.00/0.27 | — | 0.95/0.56 | 1.00/0.25 | 0.98/1.39 | 0.63/0.13 | 0.77/0.43 |
| 26826 | 1.00/0.26 | — | — | 1.00/0.24 | 0.98/1.38 | 0.58/0.13 | 0.72/0.39 |
| 26827 | 1.00/0.26 | — | — | 1.00/0.24 | 0.98/1.38 | 0.61/0.13 | 0.76/0.41 |
| 26828 | 1.00/0.30 | — | — | 1.00/0.25 | 0.98/1.39 | 0.53/0.14 | 0.63/0.44 |
| 26829 | 1.00/0.31 | — | — | 1.00/0.27 | 0.98/1.37 | 0.63/0.13 | 0.72/0.49 |
| 26830 | 1.00/0.28 | — | — | 1.00/0.27 | 0.98/1.37 | 0.63/0.13 | 0.73/0.46 |
| 26831 | 1.00/0.27 | — | — | 1.00/0.24 | 0.98/1.38 | 0.58/0.14 | 0.79/0.39 |
| 26953 | 0.99/0.87 | — | 0.88/0.73 | 1.00/0.30 | 0.96/1.31 | 0.45/0.20 | 0.41/1.09 |
| 27161 | 0.91/1.71 | — | 0.68/1.15 | 1.00/0.80 | 0.93/2.05 | 0.48/0.36 | 0.25/1.86 |
| 27290 | 1.00/0.37 | — | 0.97/0.46 | 1.00/0.31 | 0.96/1.35 | 0.55/0.15 | 0.37/0.58 |

| | | | | | | | |
|---|---|---|---|---|---|---|---|
| 27401 | 1.00/0.73 | — | 0.85/0.92 | 1.00/0.32 | 0.96/1.62 | 0.55/0.21 | 0.47/0.99 |
| 27461 | 1.00/0.67 | — | — | 1.00/0.39 | 0.94/1.36 | 0.43/0.16 | 0.41/0.61 |
| 27533 | 0.98/1.14 | — | 0.91/1.41 | 1.00/0.37 | 0.90/1.69 | 0.40/0.25 | 0.38/1.44 |
| 27569 | 0.98/1.11 | — | 0.92/0.82 | 1.00/0.35 | 0.92/1.54 | 0.48/0.23 | 0.51/1.38 |
| 27596 | 0.99/0.64 | 0.93/0.07 | 0.72/1.10 | 1.00/0.30 | 0.96/1.07 | 0.59/0.16 | 0.59/0.86 |
| 27625 | 1.00/0.61 | — | 0.91/0.53 | 1.00/0.31 | 0.95/1.71 | 0.65/0.12 | 0.22/0.54 |
| 50017 | 1.00/0.23 | — | 0.90/0.80 | 1.00/0.28 | 0.98/1.18 | 0.36/0.17 | 0.65/0.32 |
| 50113 | 1.00/0.41 | — | 0.89/0.41 | 1.00/0.46 | 0.95/1.30 | 0.67/0.14 | 0.57/0.71 |
| 50204 | 1.00/0.76 | — | 0.94/0.56 | 1.00/0.66 | 0.96/1.53 | 0.44/0.19 | 0.46/0.87 |
| 50238 | 1.00/0.28 | 0.79/0.09 | — | 1.00/0.31 | 0.96/1.20 | 0.73/0.13 | 0.54/0.48 |
| 50370 | 0.99/0.79 | — | 0.78/0.96 | 1.00/0.56 | 0.93/1.61 | 0.49/0.18 | 0.34/0.99 |
| 50542 | 0.99/0.51 | 0.93/0.13 | 0.92/0.54 | 1.00/0.28 | 0.96/1.23 | 0.58/0.15 | 0.52/0.62 |
| 50557 | 0.99/0.55 | — | 0.97/0.51 | — | 0.95/1.66 | 0.48/0.16 | — |
| 50631 | 1.00/0.62 | 0.90/0.07 | — | 1.00/0.23 | — | 0.77/0.18 | 0.43/0.71 |
| 50752 | 0.99/0.58 | 0.94/0.08 | 0.90/0.42 | 1.00/0.38 | 0.88/1.94 | 0.30/0.27 | 0.44/0.66 |
| 51011 | 1.00/0.40 | 0.97/0.06 | 0.94/0.40 | 1.00/0.25 | 0.92/1.47 | 0.48/0.18 | 0.55/0.58 |
| 51033 | 1.00/0.60 | 0.77/0.14 | 0.88/0.65 | 1.00/0.33 | 0.92/1.67 | 0.22/0.24 | 0.45/0.72 |
| 51062 | 0.99/1.16 | — | 0.94/0.42 | 1.00/1.21 | 0.95/1.47 | 0.55/0.15 | 0.30/0.68 |
| 51147 | 1.00/0.49 | 0.85/0.09 | 0.77/0.95 | 1.00/0.38 | 0.96/1.39 | 0.49/0.17 | 0.58/0.57 |
| 51354 | 1.00/0.48 | 0.96/0.06 | 0.86/0.68 | 1.00/0.29 | 0.96/1.41 | 0.48/0.18 | 0.46/0.62 |
| 51355 | 0.99/0.61 | 0.87/0.11 | 0.89/0.67 | 1.00/0.35 | 0.90/1.57 | 0.32/0.24 | 0.39/0.87 |
| 51952 | 0.99/0.51 | 0.94/0.06 | 0.91/0.60 | 1.00/0.22 | 0.94/1.14 | 0.39/0.20 | 0.74/0.66 |
| 51969 | 0.99/0.91 | — | 0.69/0.90 | 1.00/0.29 | 0.90/1.73 | 0.41/0.18 | 0.31/1.02 |
| 52005 | — | — | — | — | 0.69/3.55 | 0.20/0.68 | — |
| 52109 | 0.99/0.63 | — | 0.93/0.45 | 1.00/0.30 | 0.95/1.32 | 0.42/0.23 | 0.39/0.77 |
| 52398 | 0.99/0.45 | — | 0.90/0.50 | 1.00/0.26 | 0.92/1.40 | 0.48/0.17 | 0.51/0.58 |
| 52877 | 0.99/0.59 | 0.91/0.09 | 0.97/0.34 | 1.00/0.28 | 0.92/1.58 | 0.19/0.23 | 0.30/0.65 |
| Ntail[25] | 0.99/0.49 | — | 0.90/0.64 | 1.00/0.28 | 0.94/1.36 | 0.63/0.15 | 0.38/0.68 |

**Supplementary Table S8.** The helicity of training set containing 15 IDPs

| IDPs | Experimental helicity | HyRes helicity |
|---|---|---|
| AAQAA[18] | 0.45 | 0.44 |
| N17[19] | 0.23 | 0.26 |
| BMRB 15123 | 0.15 | 0.16 |
| BMRB 52877 | 0.11 | 0.11 |
| BMRB 15397 | 0.12 | 0.13 |
| BMRB 50238 | 0.13 | 0.14 |
| BMRB 51952 | 0.17 | 0.19 |
| BMRB 27290 | 0.08 | 0.10 |
| MevN[20] | 0.16 | 0.17 |
| BMRB 50017 | 0.04 | 0.02 |
| HevN[21] | 0.13 | 0.14 |
| BMRB 26823 | 0.08 | 0.07 |
| MevP[22] | 0.10 | 0.12 |
| BMRB 26672 | 0.05 | 0.04 |
| BMRB 50370 | 0.09 | 0.10 |

**Supplementary Table S9.** The helicity of test set containing 40 IDPs

| IDPs | Exp. helicity | HyRes helicity | IDPs | Exp. helicity | HyRes helicity |
|---|---|---|---|---|---|
| BMRB 6857 | 0.04 | 0.11 | BMRB 51355 | 0.05 | 0.09 |
| BMRB 50631 | 0.10 | 0.13 | BMRB 16798 | 0.07 | 0.07 |
| BMRB 27161 | 0.42 | 0.28 | BMRB 50542 | 0.20 | 0.13 |
| BMRB 25632 | 0.04 | 0.12 | BMRB 26762 | 0.01 | 0.07 |
| Cby[23] | 0.13 | 0.11 | BMRB 27596 | 0.23 | 0.26 |
| BMRB 51354 | 0.03 | 0.09 | BMRB 26953 | 0.09 | 0.14 |
| BMRB 50752 | 0.08 | 0.10 | BMRB 52109 | 0.05 | 0.11 |
| BMRB 27625 | 0.03 | 0.09 | BMRB 52398 | 0.07 | 0.13 |
| BMRB 26719 | 0.11 | 0.07 | BMRB 51147 | 0.07 | 0.09 |
| BMRB 18198 | 0.08 | 0.09 | BMRB 27401 | 0.04 | 0.09 |
| BMRB 15506 | 0.01 | 0.04 | BMRB 51662 | 0.09 | 0.10 |
| Sic1[24] | 0.10 | 0.07 | BMRB 25384 | 0.08 | 0.10 |
| BMRB 19830 | 0.23 | 0.19 | BMRB 51033 | 0.05 | 0.07 |
| BMRB 50557 | 0.11 | 0.13 | BMRB 50204 | 0.06 | 0.11 |
| BMRB 27461 | 0.08 | 0.10 | BMRB 26639 | 0.18 | 0.13 |
| BMRB 16165 | 0.06 | 0.13 | BMRB 52525 | 0.07 | 0.11 |
| BMRB 15141 | 0.17 | 0.19 | BMRB 26549 | 0.11 | 0.14 |
| BMRB 52005 | 0.12 | 0.16 | BMRB 51062 | 0.10 | 0.09 |
| BMRB 19332 | 0.12 | 0.10 | BMRB 51011 | 0.03 | 0.10 |
| BMRB 51969 | 0.07 | 0.12 | BMRB 50701 | 0.04 | 0.08 |

**Supplementary Table S10.** The helicity of 12 TDP-43 CTD variants

| TDP-43 CTD variants | BMRB IDs | Experimental helicity | HyRes helicity |
|---|---|---|---|
| WT | 26823 | 0.08 | 0.07 |
| A321G | 26826 | 0.07 | 0.06 |
| A321V | 26827 | 0.08 | 0.07 |
| Q331K | 26829 | 0.09 | 0.07 |
| M337V | 26830 | 0.08 | 0.07 |
| A326P | 26828 | 0.05 | 0.05 |
| M337P | 26831 | 0.07 | 0.07 |
| G335A | 27750 | 0.10 | 0.08 |
| G335N | 27788 | 0.09 | 0.08 |
| G335S | 27789 | 0.09 | 0.08 |
| G338A | 27790 | 0.10 | 0.08 |
| G335D | 50090 | 0.09 | 0.08 |

**Supplementary Table S11.** RMSD of predicted chemical shifts of 7 IDPs using HyRes, C36m, a03ws, and a99SB-disp

| Atom | Force fields | RMSD (ppm) | | | | | | |
|---|---|---|---|---|---|---|---|---|
| | | **ACTR** | **Ntail** | **aSyn** | **PaaA2** | **p15PAF** | **Sic1** | **Ash1** |
| Cα | HyRes | 0.39 | 0.49 | 0.95 | 0.74 | 0.48 | 1.02 | 0.53 |
| | C36m | 0.50 | 0.66 | 0.61 | 0.74 | 0.42 | 1.15 | 0.30 |
| | a03ws | 0.78 | 0.61 | 0.6 | 1.27 | 0.66 | 0.95 | 0.52 |
| | a99SB-disp | 0.46 | 0.44 | 0.51 | 0.64 | 0.4 | 0.77 | 0.45 |
| Hα | HyRes | — | — | — | 0.12 | 0.06 | 0.17 | — |
| | C36m | — | — | — | 0.13 | 0.09 | 0.30 | 0.15 |
| | a03ws | — | — | — | 0.17 | 0.11 | 0.26 | 0.16 |
| | a99SB-disp | — | — | — | 0.10 | 0.07 | 0.22 | 0.11 |
| HN | HyRes | 0.14 | 0.15 | 0.24 | 0.15 | 0.18 | 0.18 | 0.17 |
| | C36m | 0.21 | 0.18 | 0.2 | 0.17 | 0.14 | 0.22 | 0.52 |
| | a03ws | 0.21 | 0.18 | 0.19 | 0.22 | 0.18 | 0.17 | 0.56 |
| | a99SB-disp | 0.18 | 0.12 | 0.14 | 0.17 | 0.14 | 0.12 | 0.53 |
| N | HyRes | 1.51 | 1.36 | 1.74 | 1.34 | 1.60 | 1.52 | 1.48 |
| | C36m | 1.01 | 1.2 | 1.53 | 1.02 | 1.07 | 1.41 | 0.86 |
| | a03ws | 1.35 | 1.31 | 1.77 | 1.61 | 1.1 | 1.41 | 1.17 |
| | a99SB | 0.87 | 0.89 | 1.46 | 0.99 | 0.72 | 1.17 | 1.06 |
| C’ | HyRes | 0.42 | 0.64 | 0.58 | 0.72 | 0.57 | — | 0.74 |
| | C36m | 0.47 | 0.64 | 0.57 | 0.56 | 0.38 | — | 0.43 |
| | a03ws | 0.63 | 0.58 | 0.68 | 1.26 | 0.48 | — | 0.5 |
| | a99SB-disp | 0.42 | 0.41 | 0.31 | 0.66 | 0.36 | — | 0.41 |
| Cβ | HyRes | 0.24 | 0.28 | 0.94 | 0.37 | 0.31 | 0.52 | 0.36 |
| | C36m | 0.33 | 0.5 | 1.27 | 0.45 | 0.46 | 0.79 | — |
| | a03ws | 0.37 | 0.44 | 1.49 | 0.59 | 0.33 | 0.68 | — |
| | a99SB-disp | 0.30 | 0.37 | 1.04 | 0.37 | 0.36 | 0.52 | — |

**Supplementary Table S12.** Residue-pair distances within H1-ProTα dimer from smFRET[26] and HyRes simulations. Residues in red and blue are from ProTα and H1, respectively.

| residue 1 | residue 2 | smFRET distances (Å) | HyRes distances (Å) |
|---|---|---|---|
| 2 | 56 | 54.0 | 58.3 |
| 56 | 110 | 51.2 | 45.6 |
| 2 | 110 | 61.2 | 74.1 |
| 2 | 0 | 58.6 | 81.2 |
| 2 | 89 | 55.8 | 69.6 |
| 2 | 104 | 54.7 | 63.7 |
| 2 | 113 | 55.5 | 63.3 |
| 2 | 151 | 55.5 | 64.1 |
| 2 | 161 | 54.4 | 64.4 |
| 2 | 194 | 56.3 | 68.8 |
| 56 | 0 | 51.2 | 62.2 |
| 56 | 89 | 47.3 | 45.5 |
| 56 | 104 | 41.4 | 32.5 |
| 56 | 113 | 41.5 | 29.5 |
| 56 | 151 | 39.9 | 26.3 |
| 56 | 161 | 39.2 | 26.8 |
| 56 | 194 | 44.2 | 37.9 |
| 110 | 0 | 52.2 | 66.0 |
| 110 | 89 | 49.4 | 52.6 |
| 110 | 104 | 45.8 | 43.7 |
| 110 | 113 | 46.9 | 42.6 |
| 110 | 151 | 47.7 | 41.2 |
| 110 | 161 | 46.6 | 40.8 |
| 110 | 194 | 48.3 | 46.4 |
| 0 | 113 | 50.9 | 57.6 |
| 0 | 194 | 57.0 | 72.4 |
| 104 | 194 | 53.0 | 49.3 |
| 113 | 194 | 51.2 | 47.6 |

### Supplementary Figures

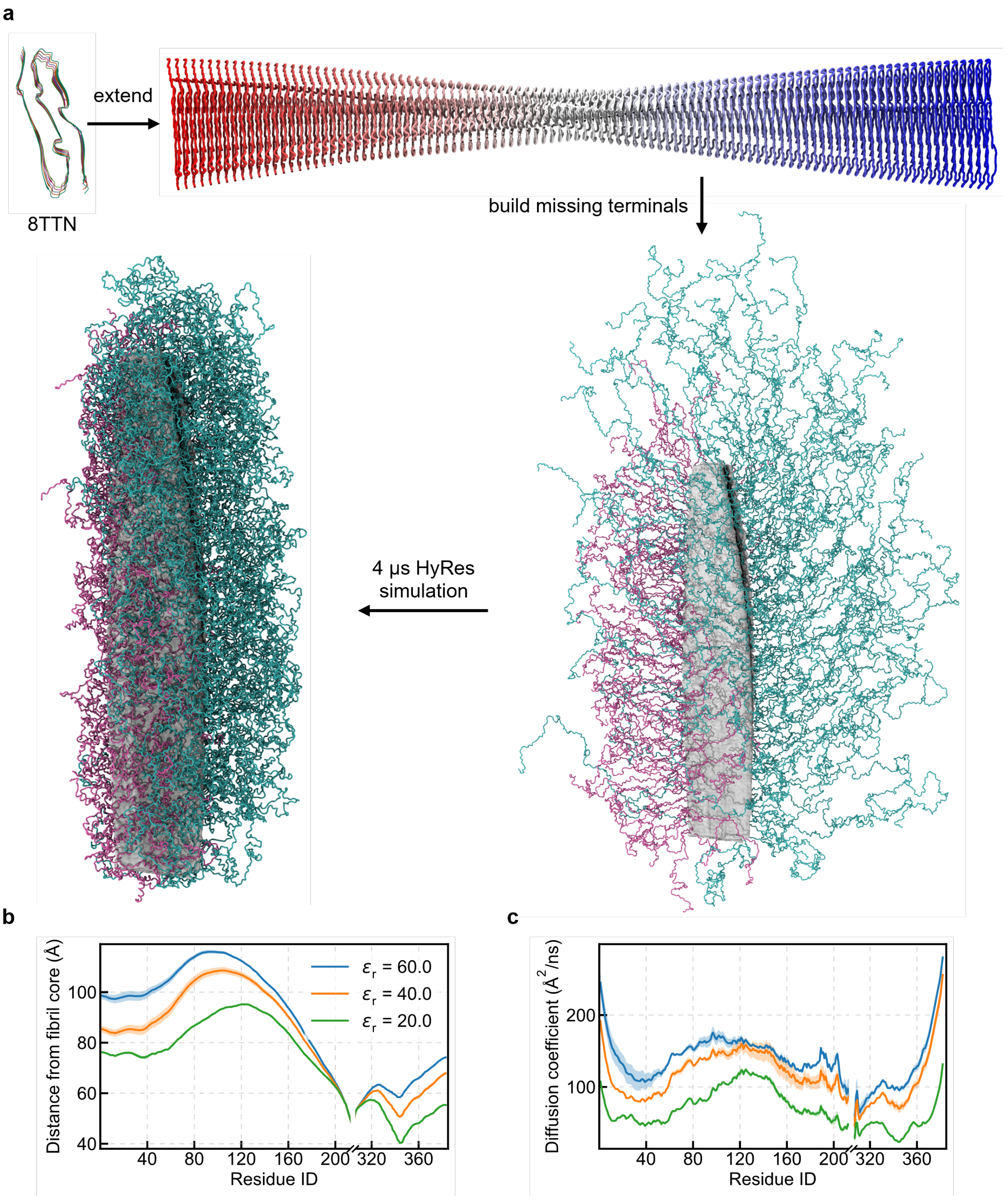


**Supplementary Fig. 1 Modeling of full-length tau fibril**. **a**, Long fibril was built from 8TTN structure through extending, missing residues modeling, and HyRes sampling. **b-c**, Radial residual distributions (**b**) and diffusion coefficients (**c**) obtained from HyRes simulations using different dielectric constant. Shadow area shows the error bar estimated as block deviation.

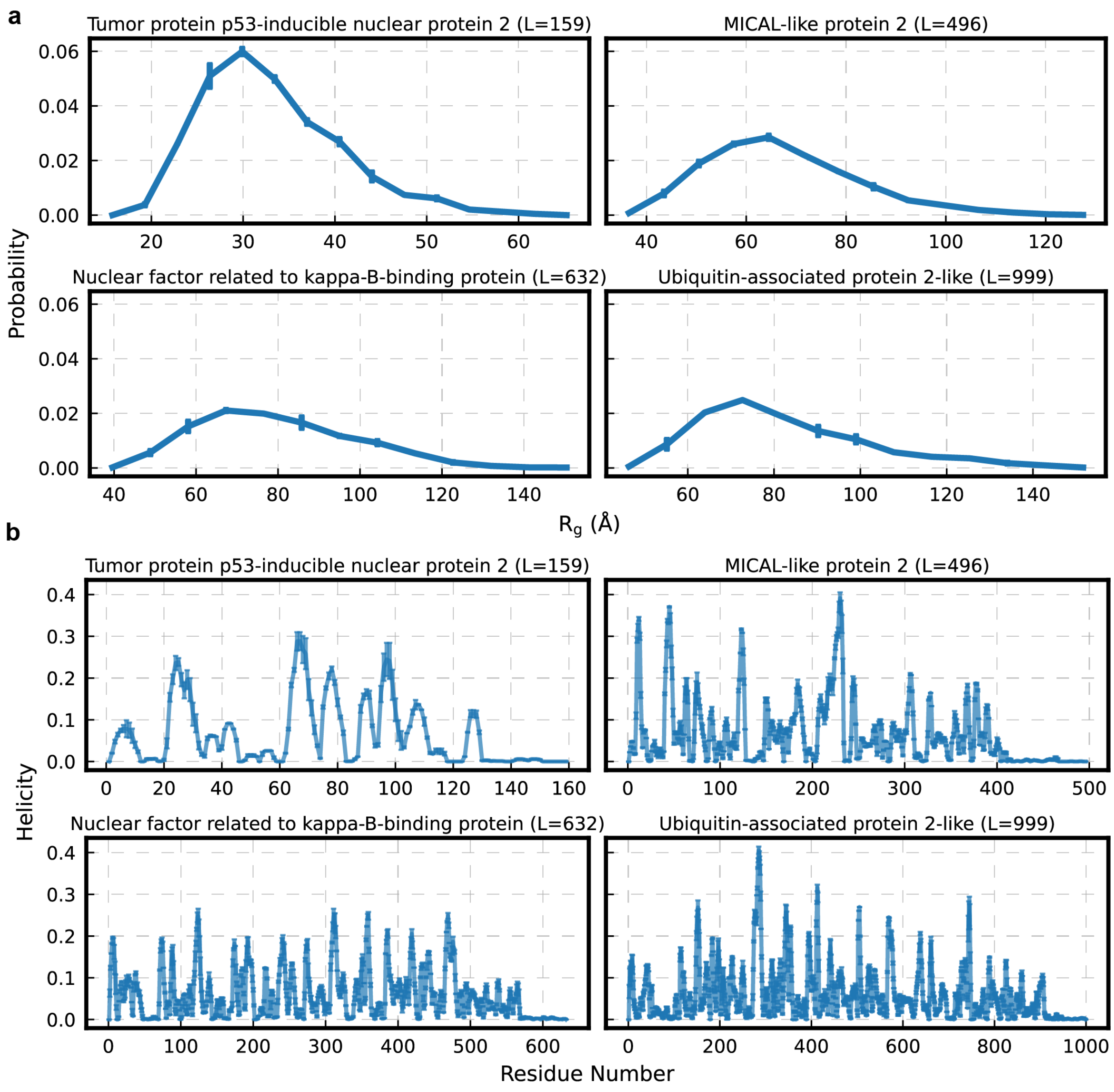


**Supplementary Fig. 2** Convergence of HyRes simulations in generation of the HyRes-IDRome database. **a-b**, $R_g$ distributions (**a**) and helicity profiles (**b**) for four sequences with different lengths. Error bars are calculated from the deviations between two halfs, indicating fully converged for both dimensional and residual structural properties.

### Supplementary References


1. Bremer, A. *et al.* Deciphering how naturally occurring sequence features impact the phase behaviours of disordered prion-like domains. *Nat. Chem.* **14**, 196–207 (2022).
2. Das, R. K. & Pappu, R. V. Conformations of intrinsically disordered proteins are influenced by linear sequence distributions of oppositely charged residues. *Proc. Natl. Acad. Sci.* **110**, 13392–13397 (2013).
3. Tesei, G., Schulze, T. K., Crehuet, R. & Lindorff-Larsen, K. Accurate model of liquid–liquid phase behavior of intrinsically disordered proteins from optimization of single-chain properties. *Proc. Natl. Acad. Sci.* **118**, e2111696118 (2021).
4. Holla, A. *et al.* Identifying Sequence Effects on Chain Dimensions of Disordered Proteins by Integrating Experiments and Simulations. *JACS Au* **4**, 4729–4743 (2024).
5. Bertoncini, C. W. *et al.* Release of long-range tertiary interactions potentiates aggregation of natively unstructured α-synuclein. *Proc. Natl. Acad. Sci.* **102**, 1430–1435 (2005).
6. Cho, M.-K. *et al.* Structural characterization of α-synuclein in an aggregation prone state. *Protein Sci.* **18**, 1840–1846 (2009).
7. Dedmon, M. M., Lindorff-Larsen, K., Christodoulou, J., Vendruscolo, M. & Dobson, C. M. Mapping Long-Range Interactions in α-Synuclein using Spin-Label NMR and Ensemble Molecular Dynamics Simulations. *J. Am. Chem. Soc.* **127**, 476–477 (2005).
8. Iešmantavičius, V. *et al.* Modulation of the Intrinsic Helix Propensity of an Intrinsically Disordered Protein Reveals Long-Range Helix–Helix Interactions. *J. Am. Chem. Soc.* **135**, 10155–10163 (2013).
9. Konrat, R. NMR contributions to structural dynamics studies of intrinsically disordered proteins. *J. Magn. Reson.* **241**, 74–85 (2014).
10. Dancheck, B., Nairn, A. C. & Peti, W. Detailed Structural Characterization of Unbound Protein Phosphatase 1 Inhibitors. *Biochemistry* **47**, 12346–12356 (2008).
11. Kubáň, V. *et al.* Quantitative Conformational Analysis of Functionally Important Electrostatic Interactions in the Intrinsically Disordered Region of Delta Subunit of Bacterial RNA Polymerase. *J. Am. Chem. Soc.* **141**, 16817–16828 (2019).
12. Du, Z. *et al.* The sequence–structure–function relationship of intrinsic ERα disorder. *Nature* **638**, 1130–1138 (2025).
13. Schwalbe, M. *et al.* Predictive Atomic Resolution Descriptions of Intrinsically Disordered hTau40 and α-Synuclein in Solution from NMR and Small Angle Scattering. *Structure* **22**, 238–249 (2014).
14. Akoury, E. *et al.* Remodeling of the conformational ensemble of the repeat domain of tau by an aggregation enhancer. *Protein Sci.* **25**, 1010–1020 (2016).
15. Lowry, D. F., Stancik, A., Shrestha, R. M. & Daughdrill, G. W. Modeling the accessible conformations of the intrinsically unstructured transactivation domain of p53. *Proteins Struct. Funct. Bioinforma.* **71**, 587–598 (2008).
16. Robustelli, P., Piana, S. & Shaw, D. E. Developing a molecular dynamics force field for both folded and disordered protein states. *Proc. Natl. Acad. Sci.* **115**, E4758–E4766 (2018).
17. Liu, X. & Chen, J. Residual Structures and Transient Long-Range Interactions of p53 Transactivation Domain: Assessment of Explicit Solvent Protein Force Fields. *J. Chem. Theory Comput.* **15**, 4708–4720 (2019).

18. Shalongo, W., Dugad, L. & Stellwagen, E. Distribution of Helicity within the Model Peptide Acetyl(AAQAA)3amide. *J. Am. Chem. Soc.* **116**, 8288–8293 (1994).
19. Urbanek, A. *et al.* Flanking Regions Determine the Structure of the Poly-Glutamine in Huntingtin through Mechanisms Common among Glutamine-Rich Human Proteins. *Structure* **28**, 733-746.e5 (2020).
20. Jensen, M. R. *et al.* Intrinsic disorder in measles virus nucleocapsids. *Proc. Natl. Acad. Sci.* **108**, 9839–9844 (2011).
21. Communie, G. *et al.* Atomic Resolution Description of the Interaction between the Nucleoprotein and Phosphoprotein of Hendra Virus. *PLOS Pathog.* **9**, e1003631 (2013).
22. Milles, S. *et al.* An ultraweak interaction in the intrinsically disordered replication machinery is essential for measles virus function. *Sci. Adv.* **4**, eaat7778 (2018).
23. Mokhtarzada, S., Yu, C., Brickenden, A. & Choy, W.-Y. Structural Characterization of Partially Disordered Human Chibby: Insights into Its Function in the Wnt-Signaling Pathway. *Biochemistry* **50**, 715–726 (2011).
24. Mittag, T. *et al.* Dynamic equilibrium engagement of a polyvalent ligand with a single-site receptor. *Proc. Natl. Acad. Sci.* **105**, 17772–17777 (2008).
25. Gely, S. *et al.* Solution structure of the C-terminal X domain of the measles virus phosphoprotein and interaction with the intrinsically disordered C-terminal domain of the nucleoprotein. *J. Mol. Recognit.* **23**, 435–447 (2010).
26. Borgia, A. *et al.* Extreme disorder in an ultrahigh-affinity protein complex. *Nature* **555**, 61–66 (2018).
